# Neuroinflammation alters the phenotype of lymphangiogenic vessels near the cribriform plate

**DOI:** 10.1101/2020.10.08.331801

**Authors:** Martin Hsu, Andy Madrid, Yun Hwa Choi, Collin Laaker, Melinda Herbath, Matyas Sandor, Zsuzsanna Fabry

## Abstract

Meningeal lymphatic vessels residing in the dural layer surrounding the dorsal regions of the brain, basal regions, and near the cribriform plate have all been implicated in the management of neuroinflammation and edema. Interestingly, only the lymphatic vessels near the cribriform plate undergo functional lymphangiogenesis in a mouse model of Multiple Sclerosis, suggesting these particular lymphatics uniquely undergo dynamic changes in response to neuroinflammation and may have distinct access to pro-lymphangiogenic factors in the CNS. However, it is unknown if these newly formed lymphangiogenic vessels are functionally similar to steady-state or if they have any other functional changes during neuroinflammation. In this study, we generated a novel protocol to isolate lymphatic endothelial cells from the cribriform plate for single cell analysis. We demonstrate that neuroinflammation-induced lymphangiogenic vessels undergo unique changes, including the capture of CNS-derived antigens, upregulation of adhesion and immune-modulatory molecules to interact with dendritic cells, and display IFN-γ dependent changes in response to the microenvironment. Single-cell trajectory analysis showed that cribriform plate lymphangiogenic vessels are post-proliferative and not generated from trans-differentiation of myeloid cells. Additionally, we show that these lymphangiogenic vessels have access to a CSF reservoir, express the water pore Aquaporin-1, and may have direct access to the CSF due to gaps in the arachnoid epithelial layer separating the dura from the subarachnoid space. These data characterize cribriform plate lymphatics and demonstrate that these vessels are dynamic structures that engage in leukocyte interactions, antigen sampling, and undergo expansion to drain excess fluid during neuroinflammation. Neuroinflammation not only induces efficient drainage of CSF but also alters the functions of lymphatic vessels near the cribriform plate.

## Introduction

Conventional lymphatic vessels within the tissue parenchyma regulate fluid homeostasis and immunity through their ability to drain fluid, cells, and antigens to the draining lymph nodes^1,2^. Intriguingly, the central nervous system (CNS) lacks conventional lymphatics within the tissue parenchyma, yet it can maintain fluid homeostasis and undergo immune surveillance^3,4,5^. The CNS maintains a relatively constant intracranial pressure despite continuous cerebrospinal fluid (CSF) production by the choroid plexus^3^, and dyes infused into the CSF drain outward and into the cervical lymph nodes^6^. Additionally, CNS derived antigens can be found in the draining lymph nodes during steady-state conditions to contribute to immunosurveillance, and drainage of CNS derived antigens increases during CNS autoimmunity^7,8,9^. Thus, how the CNS can drain fluid, cells, and antigens may lead to a better understanding of fluid dynamics, how the immune system balances immunosurveillance and autoimmunity, and how it may contribute to different brain diseases.

Historically it has been hypothesized that the majority of fluid in the subarachnoid space in animal models drain through the cribriform plate to be picked up by nearby lymphatics, which in turn drain to the cervical lymph nodes^6,9,10,11,12,13^. Many studies have infused various dyes into the CSF of animal models and visualized its exit through the cribriform plate and, subsequently, the draining lymph nodes^6^; such studies hypothesized the presence of lymphatic vessels in the olfactory submucosa with direct continuity and access to the subarachnoid space (SAS). Other groups have also demonstrated that both dorsal and basal meningeal lymphatic vessels (mLVs) can contribute to fluid uptake, suggesting multiple routes of drainage^14,15,16^. Nevertheless, both dorsal and basal mLVs seem to be separated from the SAS by an E-Cadherin^+^ arachnoid barrier^16,17^. Although basal mLVs are in closer proximity to the SAS, it is unknown how these vessels access fluid and cells in the SAS through the arachnoid barrier^16,17^.

We have previously characterized lymphatic vessels on the CNS side of the cribriform plate that dynamically undergo lymphangiogenesis in response to neuroinflammation to help facilitate the drainage of CNS-derived cells and fluid^9^. Here, we investigated if these cribriform plate lymphatic vessels (cpLVs) undergo any other phenotypic changes in addition to lymphangiogenesis using single-cell RNA sequencing (scRNAseq) during experimental autoimmune encephalomyelitis (EAE), a mouse model of Multiple Sclerosis (MS). Additionally, we compared their access to CSF with the dural and basal mLVs to determine if any unique functional changes such as lymphangiogenesis may be due to differences in fluid sampling. We find that during neuroinflammation cpLVs are functionally distinct from steady-state lymphatic endothelial cells (LECs) and upregulate genes involved in leukocyte interactions and response to the microenvironment. These expressional changes include leukocyte adhesion/chemotaxis, antigen processing and presentation, regulation of leukocyte activation, and response to interferon-gamma (IFN-γ) signaling. Functionally, cpLVs increased their ability to bind to both CD11b^+^ CD11c^high^ dendritic cells and CD4^+^ T cells, contained intracellular CNS-derived antigens, expressed MHC II, and increased the expression of several immunoregulatory proteins including the lymphocyte-tolerizing PDL-1 in an IFN-γ-dependent fashion. Additionally, cpLVs seem to have unique CSF access due to gaps in the arachnoid barrier separating the cpLVs from the subarachnoid space, increased fluid accumulation near the cribriform plate, and cpLV expression of the water pore AQP-1. These data implicate lymphangiogenic cpLVs as a potential site of immune regulation through leukocyte binding/crosstalk and as a significant route of fluid drainage during neuroinflammation.

## Results

### cpLVs undergo dynamic changes during neuroinflammation

We have previously demonstrated that neuroinflammation drives extensive VEGFR3 dependent lymphangiogenesis near the cribriform plate, which can functionally contribute to the drainage of CNS-derived antigens, cells, and fluid^9^. In order to further characterize lymphangiogenesis, we generated a single cell suspension of the cribriform plate and its associated tissues to measure cribriform plate lymphatic endothelial cell (cpLEC) numbers by flow cytometry between healthy and EAE mice **(Supplementary Figure 1A – C)**. cpLECs were identified as Podoplanin^+^ and CD31^+^ after gating for live Ghost^negative^ singlets and excluding CD45^+^ leukocytes **(Supplementary Figure 1A – B)**. Quantitation of the average number of cpLECs reveals an approximately 5-fold increase in cell number during EAE compared to healthy, validating EAE-induced expansion in LEC cell number **(Supplementary Figure 1C)**. These data confirm that cpLECs undergo expansion with increased cell numbers, and is not merely due to widening of vessels^9^.

In addition to lymphangiogenesis, we asked if cpLECs undergo any other dynamic changes in response to neuroinflammation. It is possible that lymphangiogenesis may induce other changes that make the lymphangiogenic cpLECs phenotypically unique to steady-state cpLECs. Since we have successfully developed a protocol to isolate cpLECs, we processed these cells for single-cell RNA sequencing (scRNAseq) after sorting for LECs **(Figure 1A – J)**. cpLECs were sorted as CD31^+^ Podoplanin^+^ after gating for live Ghost^negative^ singlets and excluding CD45^+^ leukocytes **(Figure 1A)**. Interestingly, both the flow analysis of LEC cell number **(Supplementary Figure 1)** and sorting for cpLECs **(Figure 1A)** seemingly show unique phenotypic differences between healthy and EAE cells, as noted particularly by the expression levels of Podoplanin, a marker commonly used to identify LECs. After sorting for cpLECs and pooling cells from 5 naïve C57BL/6J control mice separately from 5 EAE C57BL/6J mice, the cpLECs were processed with 10x Genomics Chromium Single Cell Gene Expression Assay and sequenced using the Novaseq 6000. The t-SNE plot of healthy vs. EAE cells revealed that these two populations of cpLECs separate almost exclusively into different clusters, suggesting unique phenotypes between healthy and EAE cells **(Figure 1B)**. Traditional K-means clustering grouped both healthy and EAE cpLECs into three clusters, with cluster 1 consisting primarily of healthy cells (99.84%), and clusters 2 and 3 consisting primarily of EAE cells (97.87% for cluster 2, 66.27% for cluster 3) **(Figure 1B – F)**. scRNAseq revealed that cluster 1 contained 341 differentially expressed upregulated genes (* FDR < 0.1, Methods) and 406 differentially expressed down-regulated genes; cluster 2 contained 286 differentially expressed upregulated and 393 differentially expressed down-regulated genes, and cluster 3 contained 90 differentially expressed upregulated genes with no differentially expressed down-regulated genes when locally distinguishing between the other clusters. Representative volcano plots for each of the three clusters highlight the top 50 most upregulated and downregulated genes **(Supplementary Figure 2A, C, E)**. Gene enrichment analysis of cluster 1 revealed gene pathways associated with baseline lymphatic endothelial cell function, such as catabolic process, regulation of vasculature development, and cell-cell adhesion, consistent with its makeup of primarily naive cells **(Figure 1D)**. Cluster 2, consisting primarily of EAE cpLECs, were enriched in gene pathways associated with a more activated state **(Figure 1E)** with genes involved in leukocyte crosstalk. The majority of cluster 3 consisted of EAE cpLECs (66.27%) enriched in proliferation genes, consistent with our previous data of lymphangiogenesis by cpLECs^9^ **(Figure 1F)**. Additionally, these data suggest that steady-state cpLECs undergo a baseline level of proliferation to maintain lymphatic cell numbers, and neuroinflammation seems to drive lymphangiogenesis by increasing cell proliferation.

**Figure 1:**
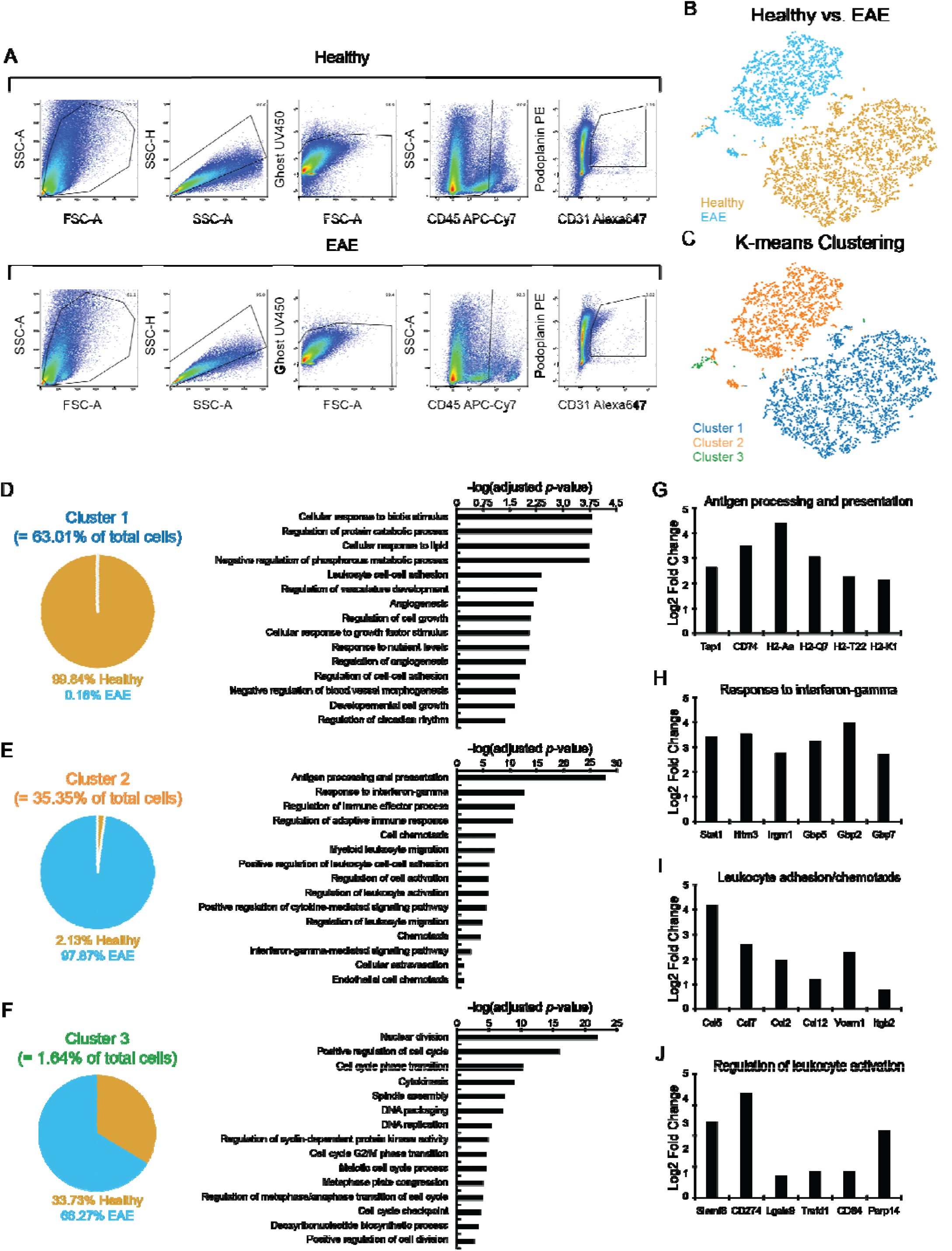
cpLVs undergo dynamic changes during neuroinflammation. **(A):** Gating strategy used to sort for cpLECs which were identified as Podoplanin^+^ CD31^+^ after excluding doublets, Ghost UV450^+^ dead cells, and CD45^+^ leukocytes. Dot plots represent a total of 5 mice per group. **(B):** t-SNE plot showing the distribution of healthy and EAE cells. **(C):** t-SNE plot showing sub-populations of cells after K-means clustering. **(D – F):** Left panels: pie chart showing the cellular composition of cluster 1 **(D)**, cluster 2 **(E)**, and cluster 3 **(F)** as shown in **(C)**. Middle panels: Representative bar plots of gene ontologies enriched for each cluster. **(G – J):** Gene ontology analysis of Cluster 2 which comprised mostly of EAE cells were sub-divided into four main categories: antigen processing and presentation **(G)**, response to IFN-γ **(H)**, leukocyte adhesion/chemotaxis **(I)**, and regulation of leukocyte activation **(J)**. Bar plots show the relative expression level of several genes within each category.

Since Cluster 2 is comprised primarily of EAE cpLECs, we decided to focus on the dynamic changes within this cluster relative to healthy cpLECs. The gene enrichment pathways were divided into four main categories: 1) antigen processing and presentation **(Figure 1G)**, 2) response to IFN-γ **(Figure 1H**, 3) leukocyte adhesion/chemotaxis **(Figure 1I)**, and 4) regulation of leukocyte activation **(Figure 1J)**, in which six representative genes associated with these pathways are shown **(Figure 1G – J)**. For each cluster, representative Cnet plots highlight the steady-state pathways in cluster 1, leukocyte crosstalk pathways in cluster 2, and proliferation pathways in cluster 3 along with all the associated genes and their relative strength of contribution to each pathway **(Supplementary Figure 2B, D, F)**. These data demonstrate that cpLECs undergo lymphangiogenic changes during neuroinflammation and may also undergo other changes in response to the environment. These changes include antigen processing/presentation, leukocyte adhesion/chemotaxis, and regulation of leukocyte activation, some of which may be in response to IFN-γ signaling as a consequence of EAE.

### Visualizing cpLEC trajectories

Since both healthy and EAE cells were almost exclusively clustered within clusters 1 and 2 with mixed cells in cluster 3 **(Figure 1)**, we next asked if a healthy cell from cluster 1 can directly transition into an EAE-like cell in cluster 2, or if it must first undergo a proliferative state like in cluster 3. Specifically, we asked if proliferation-driven lymphangiogenesis can account for all the lymphangiogenic cells in cluster 2, or if there may be some other mechanism that can account for the increased number of cpLECs during EAE. Indeed, there is evidence that inflammation-driven lymphangiogenesis may be driven through trans-differentiation of myeloid cells into LECs^18,19,20^. In line with this, we have previously demonstrated significant accumulation of myeloid cells in this region. However, we speculate these cells as not trans-differentiating but engaging in other functions such as antigen processing/presentation, adhesion/chemotaxis, and crosstalk with LECs as suggested by our scRNAseq data **(Figure 1)**. To investigate this, we visualized single-cell trajectories through pseudotime as previously described^21,22,23^, which allows us to visualize cell transitions from one state to another and consequently, can reveal how far a cell has moved through a biological process. Visualization of cell trajectories through pseudotime revealed that a steady-state cell in cluster 1 seems to transition into a proliferative state in cluster 3 on its way to become an EAE-like state in cluster 2 **(Supplementary Figure 3A)**. Visualizing specific genes revealed an upregulation of adhesion/chemotaxis genes such as CCL12, CCL2, CCL5, and CCL7 **(Supplementary Figure 3B)**; response to IFN-γ genes such as Ifitm3, Gbp2, and Stat1 **(Supplementary Figure 3C)**; and antigen processing and presentation genes such as H2-Aa, H2-Q7, and Tap1 **(Supplementary Figure 3D)** through pseudotime relatively early, while genes involved in leukocyte activation such as Parp14, Slamf8, and CD274 (PDL-1) **(Supplementary Figure 3E)** seemed to be upregulated slightly later through pseudotime. These data suggest that lymphangiogenesis through the proliferation of pre-existing LECs seems to drive many phenotypic changes of cpLECs during EAE. Additionally, the proliferation of pre-existing lymphangiogenic vessels seem to account for all the EAE-like cells in cluster 2, excluding the possibility of a direct transition of a “steady-state” cell from cluster 1 to an “EAE-like” cluster 2 cell. Additionally, we could not find any unique trace myeloid lineage genes as evident by the lack of a unique cluster that would potentially give rise to the “EAE-like” cells in cluster 2, ruling out the possibility of transdifferentiation by myeloid cells.

### Neuroinflammation promotes dendritic cell – lymphatic endothelial cell binding

Functionally, lymphatic vessels can regulate immunity by facilitating leukocyte trafficking^1,2^ and even direct regulation through ligands such as PDL-1^24,25,26,27^ or antigen presentation^26,27,28^. Indeed, scRNAseq data reveal an enrichment in antigen processing/presentation, leukocyte adhesion/chemotaxis, and regulation of leukocyte activation, including CD274 (PDL-1) **(Figure 1)**, all of which indicate binding and interaction between cpLECs and leukocytes. Therefore, we looked to see if EAE functionally promotes leukocyte binding to cpLECs, and if so, which leukocyte subpopulations are involved in leukocyte – LEC binding. Using flow cytometry, we reasoned that a doublet that expresses both LEC markers and leukocyte markers should functionally consist of a LEC bound to a leukocyte, which has previously been characterized as a method to study cell-cell interactions^29,30^. We compared healthy and EAE cell interactions by gating on live doublets that express both the pan-leukocyte CD45 marker and cpLEC markers CD31, Podoplanin, and Lyve-1 **(Figure 2A)**. Dendritic cells were defined as CD45^hi^, CD11b^+^, CD11c^high^; macrophages as CD45^hi^, CD11b^+^, CD11c^low^; CD4 T cells as CD45^high^ CD11b^-^ CD11c^-^ CD4^+^, CD8 T cells as CD45^high^ CD11b^-^ CD11c^-^ CD8^+^, and B cells as CD45^hi^ CD11b^-^ CD11c^-^ CD4^-^ CD8^-^ B220^+^. Consistent with an expansion of cpLVs during EAE, there was an approximately 50-fold increase in cpLEC cell number bound to CD45^+^ leukocytes, suggesting an increased affinity for cell binding by cpLECs during EAE **(Figure 2B)**. Of the leukocytes, a significant increase in the number of CD11b^+^ CD11c^high^ dendritic cells (≈ 150-fold higher than steady-state) and to a lesser extent CD4^+^ T cells (≈ 15-fold higher than steady-state) were bound to cpLECs, with a trending but non-significant increase in cell number of CD11b^+^ CD11c^low^ macrophages **(Figure 2B)**. This data is consistent with dendritic cells’ role as professional antigen-presenting cells that can migrate to lymphatics from the tissue parenchyma^31^ and validates our previous study demonstrating a large accumulation of inflammatory dendritic cells near the cribriform plate to contribute to VEGFC – VEGFR3 dependent lymphangiogenesis of cpLECs^9^. No elevation in CD8^+^ T cells or B220^+^ B cells were bound to cpLECs during EAE **(Figure 2B)**. Of the leukocytes bound to cpLECs, CD11b^+^ CD11c^+^ dendritic cells made up the majority of cells bound, again consistent with their role as professional antigen-presenting cells that traffic to the lymph node through lymphatics **(Figure 2C)**. We excluded the possibility of blood-derived leukocytes binding to cpLECs either *in vivo* through direct access between blood and lymphatic vessels, or *ex vivo* through blood-contamination by intravenously (IV) labeling blood-derived leukocytes with CD45.2 conjugated to BV711 *in vivo* 3 minutes before harvest **(Supplementary Figure 4)**. After perfusion, the majority of leukocytes binding to cpLECs were not blood-derived (≈ 96%) **(Supplementary Figure 4)**, suggesting that the leukocytes binding to cpLECs came from the tissue parenchyma. In order to further validate our flow cytometry data, we induced EAE in CD11c-eYFP transgenic reporter mice and harvested cpLECs for visualization with confocal microscopy using the same harvest process for flow cytometry and stained for cpLECs using CD31 and Podoplanin. Doublets in the cell suspension consisted primarily of a CD11c-eYFP^+^ cell bound to a CD31^+^ Podoplanin^+^ cell **(Figure 2D)**.

**Figure 2:**
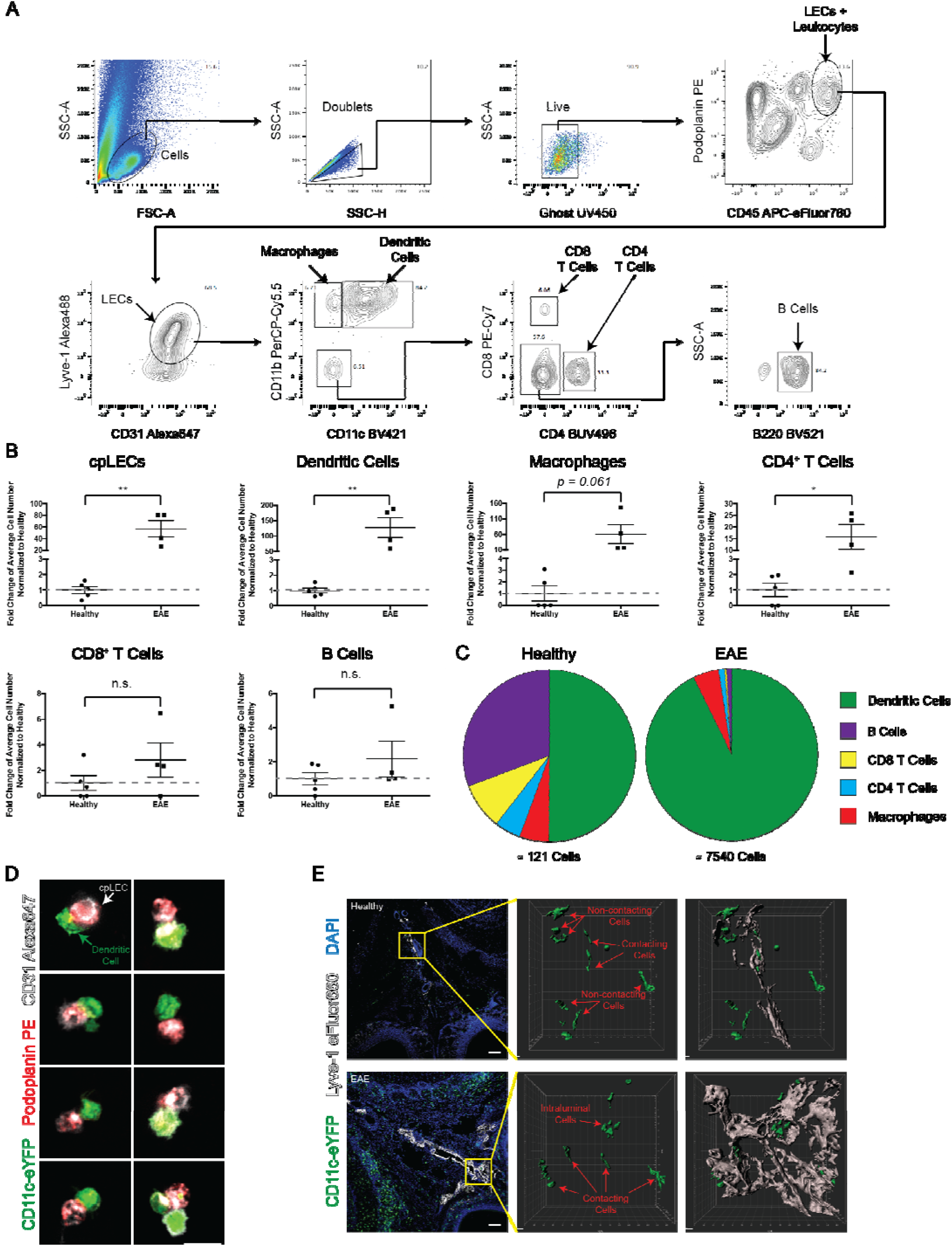
Neuroinflammation promotes cpLEC – leukocyte interactions. **(A):** Gating strategy used to visualize leukocyte – LEC binding. Live Ghost UV450^-^ doublets were gated for, and a leukocyte bound to a LEC were gated for both the leukocyte marker CD45 and LEC markers Podoplanin, Lyve-1, and CD31. Leukocytes were further gated for CD11b^+^ macrophages, CD11b^+^ CD11c^+^ dendritic cells, CD4^+^ T cells, CD8^+^ T cells, and B220^+^ B cells. **(B):** Quantitation of the average fold changes in cpLECs, dendritic cells, macrophages, CD4 and CD8 T cells, and B cell numbers during EAE relative to healthy. *n* = 4 – 5 mice per group; data are represented as mean ± standard error of the mean, \**p* ≤ 0.05, \*\**p* ≤ 0.01, unpaired Student’s t-test. **(C):** Pie charts showing the relative composition of different leukocytes bound to cpLECs in healthy controls and EAE mice. **(D):** CD11c-eYFP transgenic reporter mice were induced with EAE, and underwent the same harvest protocol as **(A – C)** to visualize leukocyte – LEC binding using confocal microscopy. Representative confocal microscopy images show doublets and the occasional triplet consisting primarily of a CD11c^+^ dendritic cell bound to a CD31^+^ Podoplanin^+^ LEC. **(E):** CD11c-eYFP transgenic reporter mice were induced for EAE, and dendritic cell – LEC interactions were visualized *in situ* using immunohistochemistry. The top panel is representative of healthy, and the bottom panel is representative of EAE animals.

In some cases, CD31 was expressed by both CD11c-eYFP^+^ cells and Podoplanin^+^ LECs **(Figure 2D)**, consistent with its function as a homophilic adhesion molecule^25^. We then validated our cpLEC – leukocyte cell-binding data using coronal sections of EAE CD11c-eYFP transgenic reporter mice. IMARIS 3D surface volume rendering identified several CD11c-eYFP^+^ cells in contact with cpLECs during EAE **(Figure 2E)**, consistent with our previous study^9^. These data confirm that in response to neuroinflammation, CD11b^+^ CD11c^high^ dendritic cells and, to a lesser extent, CD4^+^ T cells can functionally bind and interact with cpLECs during EAE.

### cpLECs can capture CNS-derived antigens and express MHC II

Intriguingly, lymphatic endothelial cells in the lymph nodes have been shown to capture and archive antigens for relatively long periods, and consequently engage in antigen presentation^26,28,32^. Indeed, scRNAseq data suggest that cpLECs can engage in antigen processing and presentation through the upregulation of MHC proteins, and flow cytometry suggests an increased number of primarily CD11b^+^ CD11c^high^ dendritic cells and some CD4^+^ T cells binding to cpLECs during EAE, suggesting potential cross-talk. Therefore, we next looked to see if we could find traces of CNS-derived antigen capture by cpLECs during EAE as well as confirm their expression of MHC. We induced EAE in CNP-OVA mice, in which OVA-GFP is endogenously expressed under the CNS oligodendrocyte CNPase promoter as previously described^7,9^. These transgenic mice express OVA-GFP by oligodendrocytes, which can functionally contribute to the proliferation of OVA-specific T cells in the draining lymph nodes during neuroinflammation^7,9^. Importantly, these mice allowed us to investigate if cpLVs can gain access to CNS-derived antigens without disrupting the arachnoid barrier through cisterna magna or intracerebral injection of the exogenous peptide. Immunolabeling of the cortex in CNP-OVA transgenic mice with antibodies against GFP and CNPase confirms OVA-GFP expression within oligodendrocytes of the CNS **(Figure 3A – C)**. During EAE, OVA-GFP^+^ expression could be observed near and within Podoplanin^+^ MHC II^+^ cpLECs **(Figure 3D – H, yellow arrowheads)**, suggesting that cpLVs have access to CNS-derived antigens and can, in fact, express MHC II. Plot profile intensity analysis of cpLECs and nearby cells from panel H reveal that not all cpLECs contain MHC II **(Figure 3D – H, red arrowheads)**, and other cells nearby can also express MHC II or contain OVA-GFP **(Figure 3I)**, suggesting heterogeneity between LECs. Immunolabeling for CD11c^+^ MHC II^+^ dendritic cells confirmed their presence near Podoplanin^+^ cpLECs as well as their capability of containing OVA-GFP **(Figure 3J – N, yellow arrowheads)**, consistent with their role as professional antigen-presenting cells^31^.

**Figure 3:**
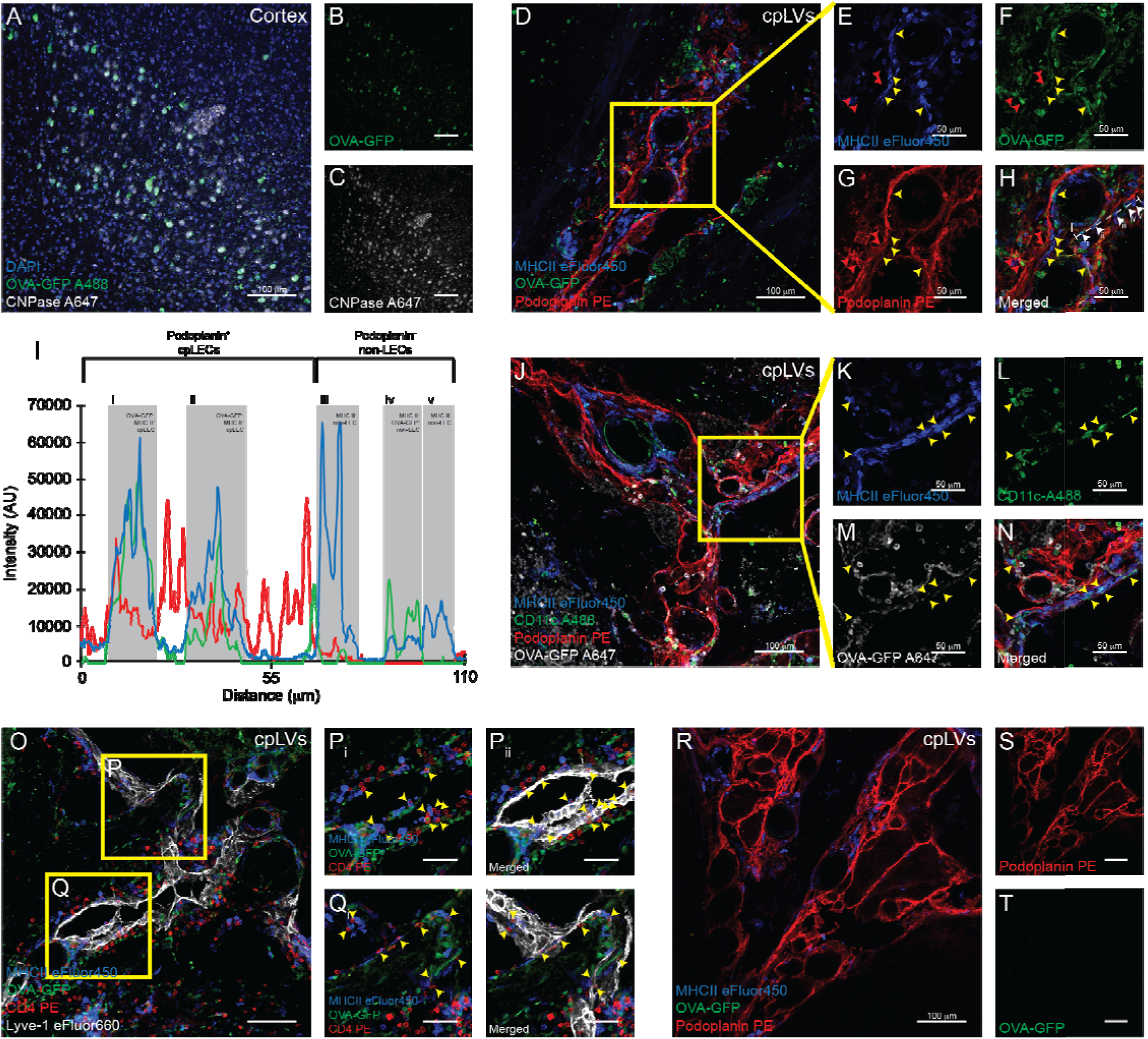
cpLECs contain CNS-derived antigens and express MHC II. **(A – C):** EAE was induced in CNP-OVA transgenic mice, in which OVA-GFP^+ fl/fl^ expression is driven by the oligodendrocyte specific CNPase-Cre. Immunohistochemistry of layers V/VI of the cortex confirm OVA-GFP expression by CNPase^+^ oligodendrocytes within the CNS when immunolabeled with an anti-GFP antibody. **(D – H):** Immunolabeling of cpLVs confirm MHC II expression and OVA-GFP^+^ signal by a subset of Podoplanin^+^ cpLECs. Red arrowheads indicate MHC II^-^ OVA-GFP^+^ cpLECs, and yellow arrowheads indicate MHC II^+^ OVA-GFP^+^ cpLECs. **(I):** Representative plot profile intensity of cells as indicated by the white line shown in **(H)** showing 5 representative cells, consisting of MHC II^+^ OVA-GFP^+^ cpLECs (i and ii), MHC II^+^ non-cpLECs (iii and v), and MHC II^+^ OVA-GFP^+^ non-cpLEC (iv). **(J – N):** Immunolabeling of cpLVs with Podoplanin and dendritic cells near the cribriform plate with CD11c along with MHC II and OVA-GFP. Yellow arrowheads indicate MHC II^+^ OVA-GFP^+^ CD11c^+^ dendritic cells in contact with Podoplanin^+^ cpLECs. **(O – Q_ii_):** Immunolabeling of cpLVs with Lyve-1 and CD4^+^ T cells near the cribriform plate along with MHC II and OVA-GFP. Yellow arrowheads indicate CD4^+^ T cells in contact with MHC II^+^ OVA-GFP^+^ Lyve-1^+^ cpLECs. **(R – T):** Control immunolabeling of Podoplanin^+^ cpLECs with MHCII and without an anti-GFP primary antibody, confirming that there is no unspecific labeling of GFP with the secondary antibody.

Similarly, CD4^+^ T cells could also be observed in this region, many of which seem to be in close contact with MHC II^+^ cpLECs and MHC II^+^ dendritic cells **(Figure 3O – Q_ii_, yellow arrowheads)**, suggesting potential crosstalk between cpLECs, dendritic cells, and CD4^+^ T cells. Control sections immunolabeled with Podoplanin, MHC II, and secondary antibody against GFP without GFP primary antibody confirmed OVA-GFP expression in this region is not due to unspecific secondary antibody binding or background **(Figure 3R – T)**. These data validate our scRNAseq data and highlight the capability of cpLECs to access CNS-derived antigen, express MHC II, and potentially engage in crosstalk with dendritic cells and CD4 T cells **(Figure 1)**. Additionally, the accumulation of both MHC II^+^ CD11c^+^ dendritic cells containing CNS-derived antigen and CD4^+^ T cells with cpLECs confirms our doublet flow data showing cpLEC – dendritic cell and cpLEC – CD4 T cell binding **(Figure 2)**, further highlighting potential crosstalk between the three cell types. While uptake of antigens and antigen presentation through MHC II have previously been attributed to lymphatic endothelial cells in the lymph nodes^20,22^, here we show that cpLECs upstream of the CNS-draining lymph nodes may also potentially process antigen and engage in antigen presentation through MHC II.

### Neuroinflammation alters cpLV expression of CD31, Podoplanin, Lyve-1, and PDL-1

While characterizing cpLVs by flow cytometry, we noticed that many markers used to identify lymphatic endothelial cells were upregulated at the protein level during neuroinflammation. Many of these markers, including Podoplanin, Lyve-1, and CD31, are all implicated in the lymphatic vessel – leukocyte interactions^33,34,35,36^, consistent with our previous data showing increased cpLEC – leukocyte binding **(Figure 2)**. While scRNAseq was unable to screen for these particular markers with sufficient average counts, other genes such as CD274 (PDL-1), a tolerogenic ligand that can interact with the PD-1 receptor, expressed by dendritic cells and T cells during inflammation^24,25,26,27^, was enriched by cpLECs during EAE **(Figure 1)**. In order to investigate if these same markers correlated with dendritic cell and CD4 T cell binding to cpLECs, we validated Podoplanin, Lyve-1, CD31, and PDL-1 upregulation by cpLVs during EAE at the protein level using flow cytometry **(Supplementary Figure 5)**. When gating for Ghost UV450 negative, singlet cpLECs **(Supplementary Figure 5A)**, the expression of Lyve-1 **(Supplementary Figure 5B – E)**, CD31-1 **(Supplementary Figure 5F – I)**, Podoplanin **(Supplementary Figure 5J – M)**, and PDL-1 **(Supplementary Figure 5N – Q)** were upregulated during EAE compared to healthy controls in the singlet gates. The expression of these proteins was further upregulated when gating for cpLEC – leukocyte doublets with the exception of Lyve-1, which may reflect that leukocytes preferably bind to cpLECs with the highest expression of these proteins. Alternatively, CD31^33^ and PDL-1^25^ have also been reported to be expressed by leukocytes/dendritic cells, respectively, which may increase the doublet median fluorescence intensity (MFI) when bound to a cpLEC that also expresses these markers. Increased background MFI when gating for doublets versus singlets could be observed for some markers, but the increase due to doublet background was negligible compared to actual protein expression **(Supplementary Figure 6)**. Increased cpLEC expression of Podoplanin, Lyve-1, CD31, and PDL-1 during EAE suggests that cpLECs alter their phenotype at the protein level to potentially promote leukocyte adhesion/chemotaxis and crosstalk through Podoplanin, Lyve-1, CD31, and PDL-1.

### cpLV upregulation of Podoplanin and PDL-1 during EAE is mediated by IFN-γ

EAE induced in C57BL/6 mice is characterized by a robust Th1/Th17 response^37^. Additionally, IFN-γ has been hypothesized to regulate Podoplanin and PDL-1 expression on lymphatic vessels during peripheral inflammation^38,39^. Coincidentally, our scRNAseq data shows enrichment in IFN-γ response genes by cpLECs during EAE, which includes CD274 (PDL-1) **(Figure 1)**. Of note, mice deficient in the pro-inflammatory IFN-γ cytokine surprisingly have exacerbated EAE^40^, suggesting a tolerogenic role of IFN-γ, which would coincide with its regulation of leukocyte activation through ligands such as PDL-1. Thus, we hypothesized that IFN-γ might be responsible for the increased expression of Podoplanin and PDL-1 by cpLECs, which may partially explain the tolerogenic role of IFN-γ during EAE. Wild-type and IFN-γ knockout mice were induced with EAE, and at EAE score 3.0, both strains were analyzed for the expression of Podoplanin and PDL-1 **(Figure 4A – I)**. Identically scored mice were used to prevent any effects that might be a consequence of differences in EAE severity. Similar to our previous results, EAE upregulated Podoplanin, CD31, and PDL-1 expression by cpLECs, as well as PDL-1 expression by dendritic cells **(Figure 4B – I)**. IFN-γ KO mice have reduced cpLEC expression of Podoplanin and PDL-1 while CD31 expression was unchanged, suggesting Podoplanin and PDL-1 upregulation during EAE are both dependent on IFN-γ signaling **(Figure 4B – I)**.

**Figure 4:**
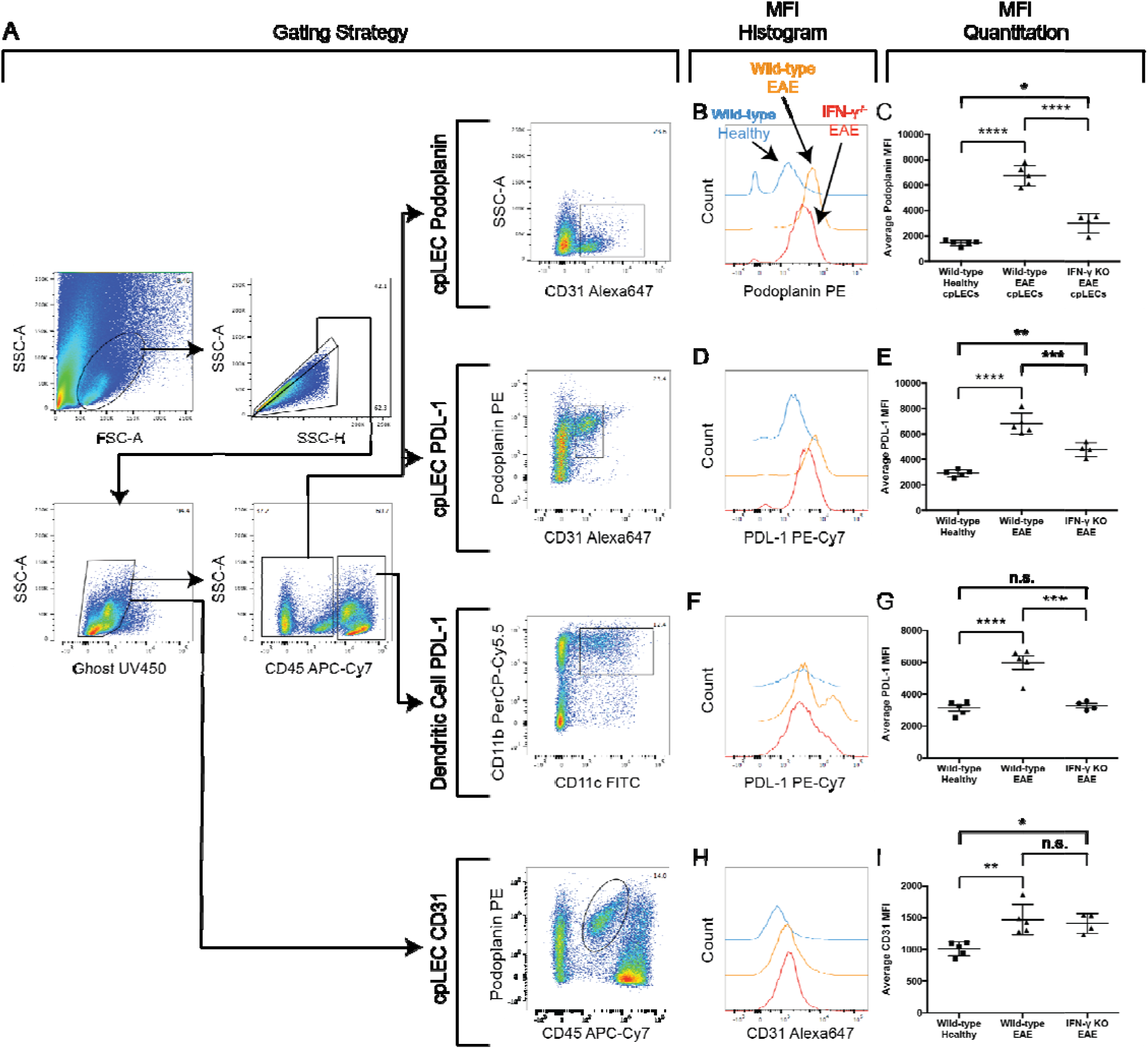
IFN-γ regulates cpLV expression of PDPN and PDL-1. **(A):** Representative gating strategy used to characterize IFN-γ dependent regulation of Podoplanin and PDL-1. EAE was induced in wild-type and IFN-γ^-/-^ C57BL/6J mice, and the expression of IFN-γ mediated Podoplanin, PDL-1, and CD31 was analyzed by flow cytometry. Gating strategy taken from a representative wild-type EAE sample. **(B – I):** The median fluorescence intensity of Podoplanin by cpLECs **(B – C)**, PDL-1 by cpLECs **(D – E)**, PDL-1 by dendritic cells **(F – G)**, and CD31 by cpLECs **(H – I)** were quantified. *n* = 4 – 5 mice per group; data are represented as mean ± standard error of the mean, \**p* ≤ 0.05, \*\**p* ≤0.01, \*\*\**p* ≤0.001, \*\*\*\**p* ≤0.001, one-way ANOVA with Tukeys post-hoc multiple comparisons test.

Interestingly, cpLECs in the IFN-γ KO mice still underwent lymphangiogenesis **(Supplementary Figure 7)**, suggesting lymphangiogenesis is not dependent on IFN-γ signaling. These data suggest that IFN-γ plays both a pro-inflammatory and tolerogenic role in EAE^40^, potentially through driving cpLV expression of PDL-1, which may partially explain the exacerbated EAE found in IFN-γ deficient mice. Additionally, other drivers of lymphangiogenesis, such as Type I interferons, have been shown to regulate some aspects of the LEC phenotype in the lymph nodes^25^.

### Lymphangiogenic vessels near the cribriform plate are in a superior position to sample CSF

In addition to leukocyte trafficking and cross-talk, another major function of lymphatic vessels is to facilitate the drainage of fluid^1,2,3^. We and others have previously demonstrated that dorsal mLVs, basal mLVs, and cpLVs can drain CSF^9,14,15,16^, yet how these lymphatics gain access through the arachnoid barrier is unknown^17^. The observation that cpLECs uniquely undergo lymphangiogenesis during EAE suggests these particular lymphatics may have distinct access to the pro-lymphangiogenic ligand VEGFC. To visualize the epithelial cells that make up the arachnoid barrier separating the dorsal mLVs, basal mLVs, and cpLVs from the subarachnoid space, we immunolabeled coronal sections with the epithelial adherence junction protein E-Cadherin^16^ **(Figure 5)**. Our data validate Ahn et al.’s studies in which there is a continuous, uninterrupted E-Cadherin^+^ arachnoid barrier separating dorsal and basal mLVs from the subarachnoid space, with basal mLVs residing closer in proximity to the subarachnoid space^16^ **(Figure 5B – C)**. Like Weller et al.’s study, there are several significant gaps in E-Cadherin labeling near the cribriform plate^17^ **(Figure 5A)**. Interestingly, here we show that the gaps in E-cadherin expression by the arachnoid barrier near the cribriform plate coincide specifically with the presence of lymphatics; there seems to be smooth, uninterrupted E-Cadherin staining where there is a lack of cpLVs and significant gaps in E-Cadherin staining where cpLVs are present **(Figure 5A)**. These data suggest that lymphatics near the cribriform plate may have greater or even direct access to fluid relative to both basal and dorsal mLVs.

**Figure 5:**
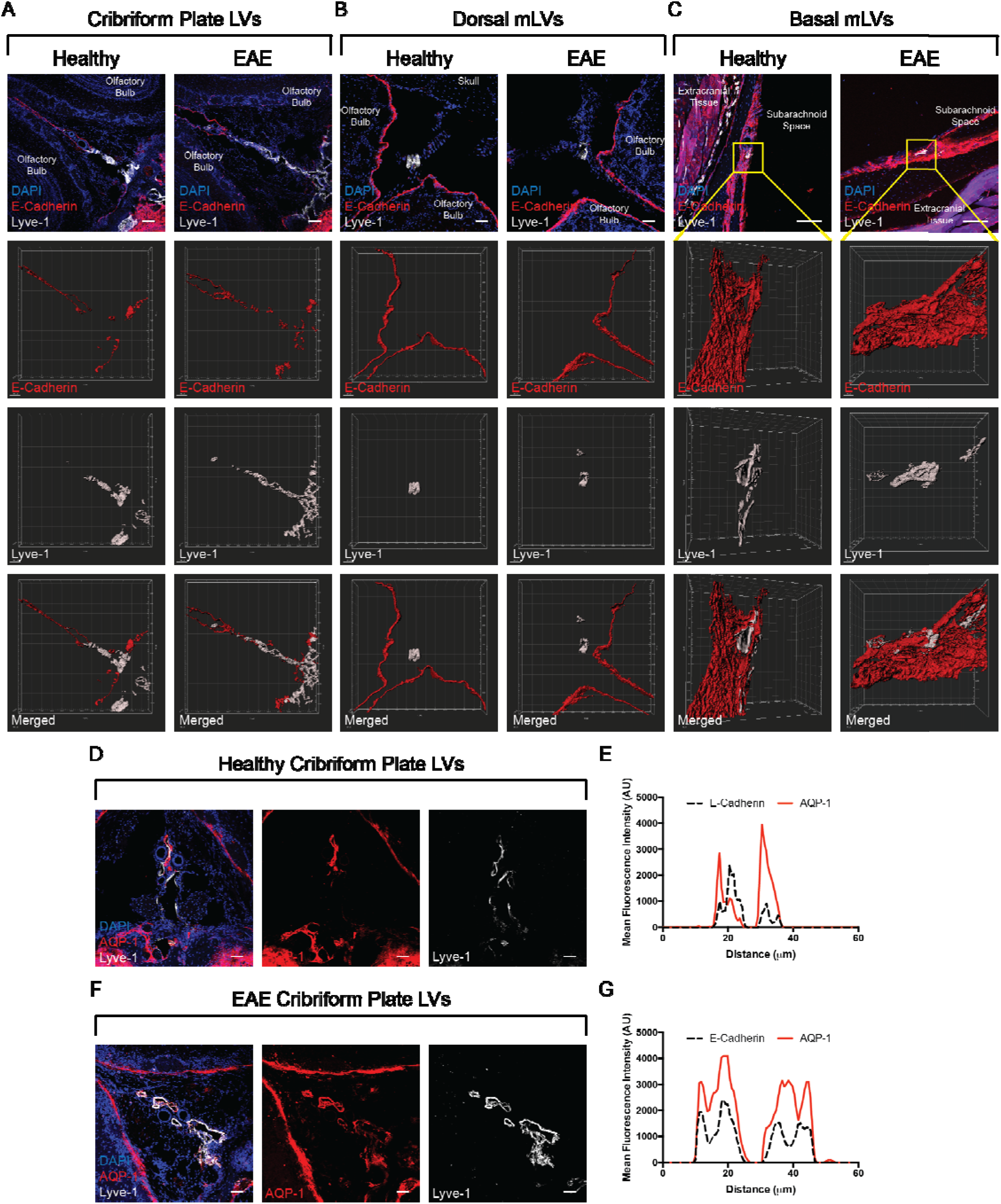
cpLVs are in a prime position to sample CSF. **(A – C):** Immunolabeling of the cribriform plate **(A)**, dura above the olfactory bulbs **(B)**, and basal meninges below the brain **(C)** from coronal sections of the whole-head. Top row: Merged immunohistochemistry images labeled with DAPI to visualize nuclei, E-Cadherin for the epithelial cells that comprise the arachnoid barrier, and Lyve-1 for lymphatic vessels. Second row: 3D surface rendering of the E-Cadherin^+^ arachnoid barrier. Third row: 3D surface rendering of the Lyve-1^+^ lymphatic vessels. Fourth row: 3D surface rendering of both the E-Cadherin^+^ arachnoid barrier and Lyve-1^+^ lymphatic vessels together. **(D – G):** Immunolabeling of the cribriform plate from either healthy **(D)** or EAE **(F)** for the water channel AQP-1 and Lyve-1. Representative plot profile intensity analysis confirms co-localization of AQP-1 expression with Lyve-1^+^ lymphatic vessels in both healthy **(E)** and EAE **(G)**.

Since we have observed that cpLECs may anatomically have greater access to the subarachnoid space than the other lymphatics, we next looked to see if they express any water transport proteins that might facilitate sampling of CSF. Aquaporins (AQPs) are often implicated in regulating fluid dynamics, with AQP-1 and AQP-4 most highly expressed in the CNS^41,42^. AQP-4 is expressed by astrocytic endfeet, hypothesized to regulate glymphatic function^43,44,45^, while AQP-1 is expressed on the apical membrane of the choroid plexus and is believed to be involved in CSF production^43^. Interestingly, a recent study by Norwood et al. reported expression of AQP-1 near the cribriform plate, although the identity of the cell types that expressed AQP-1 was unknown^13^. Here, we reveal the expression of AQP-1 by cpLVs, highlighting a potential mechanism of fluid transport from the subarachnoid space into cpLVs **(Figure 5D – G)**. Additionally, AQP-1 is expressed throughout the cpLECs, even after the lymphangiogenic expansion of cpLECs during EAE **(Figure 5D, F)**. These data suggest that there may be compensatory mechanisms to manage neuroinflammation-induced edema with increased AQP-1 expression by lymphangiogenic cpLVs. Of note, AQP-1 expression was also observed by the glia limitans surrounding the olfactory bulbs **(Figure 5D, F)**. The gaps in the arachnoid barrier and expression of the water pore AQP-1 by cpLVs highlight their increased access to CSF relative to the dorsal and basal mLVs, which may explain their unique ability to undergo lymphangiogenesis.

### cpLVs have access to a CSF reservoir

Fluid accumulation due to inflammation in the CNS may be life-threatening due to the rigidity of the skull^37^. Since we have shown that cpLVs may have direct access to CSF and undergo lymphangiogenesis to potentially increase fluid drainage during neuroinflammation^9^ **(Figure 5)**, we therefore hypothesize that CSF should also accumulate near the cribriform plate during steady-state conditions and increase its accumulation during EAE. In other words, we hypothesize that cpLVs should have access to a CSF reservoir. To visualize CSF dynamics, we employed Magnetic Resonance Imaging (MRI) before and after Gadolinium injection into the cisterna magna **(Figure 6A – E)**. Similar to a previous report, most of the Gadolinium accumulation over time occurred at the base of the brain near basal mLVs^16^ **(Figure 6B)**. However, we also discovered a relatively equal amount of Gadolinium accumulation near the base of the olfactory bulbs near cpLVs, suggesting that cpLVs have access to a reservoir of CSF **(Figure 6B)**. Additionally, our data revealed a relative increase in Gadolinium accumulation near the base of the brain and cribriform plate during EAE, consistent with the inflammation-induced edema **(Figure 6B, D)**. Consequently, there was an increase in Gadolinium accumulation in the deep cervical lymph nodes **(Figure 6C, E)**, reflecting increased fluid drainage surrounding the CNS. This data not only suggest cpLVs have access to a CSF reservoir, but that lymphangiogenesis potentially occurs in response to neuroinflammation to manage edema.

**Figure 6:**
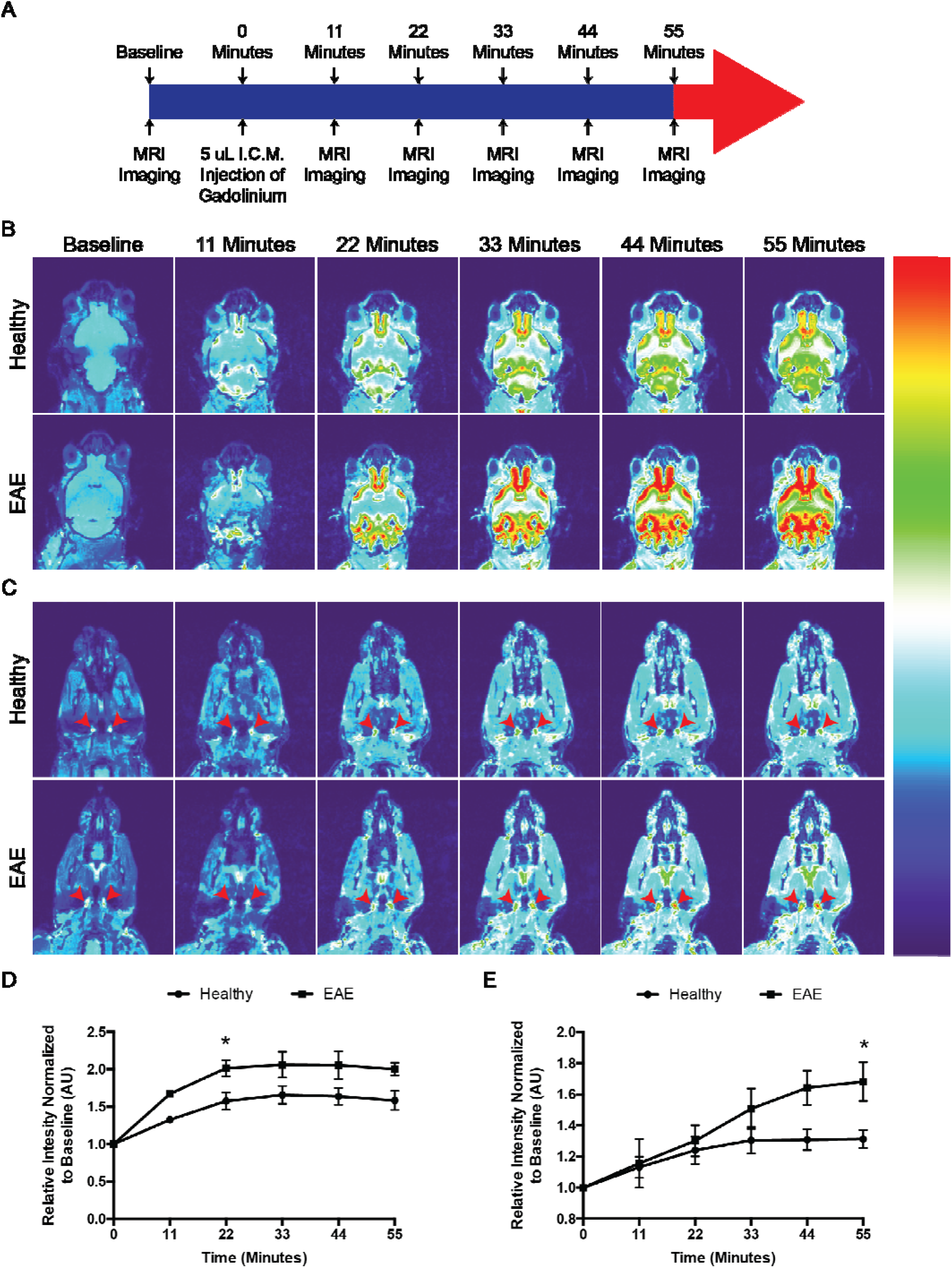
cpLVs have access to a CSF reservoir. **(A):** Schematic of experimental design. **(B):** Representative T1-weighted scans of the base of the brain and olfactory bulbs where the basal mLVs and cpLVs reside at baseline before Gadolinium infusion, and serial T1-weighted scans after Gadolinium infusion. Note the accumulation of Gadolinium signal between the olfactory bulbs and base of the brain. **(C):** Representative scans of the neck showing the dCLNs as indicated by the red arrowheads. **(D):** Quantitation of the average pixel intensity where cpLVs reside, ventral medially to the olfactory bulbs normalized to baseline between healthy and EAE. *n* = 4 mice per group; data are represented as mean ± standard error of the mean, \**p* ≤ 0.05, repeated two-way ANOVA using Sidaks multiple comparisons test. **(E):** Quantitation of the average pixel intensity of the dCLNs as indicated by the red arrowheads in **(C)** normalized to baseline between healthy and EAE. *n* = 4 mice per group; data are represented as mean ± standard error of the mean, \**p* ≤ 0.05, repeated two-way ANOVA using Sidaks multiple comparisons test.

## Discussion

Here, we provide a detailed characterization showing that cpLECs undergo several phenotypic changes in addition to lymphangiogenesis during neuroinflammation. These changes include the upregulation of genes involved in leukocyte cross-talk and response to the microenvironment. Additionally, we identify anatomical heterogeneity between the dorsal mLVs, basal mLVs, and cpLVs in terms of their access to CSF. Indeed, studies by other groups have also described heterogeneity between lymphatics. Compared to peripheral lymphatics, RNA sequencing data has shown the dorsal mLVs to be dysregulated in several aspects such as development and proliferation, extracellular matrix, focal adhesion, angiogenesis, etc^38^. Developmental studies have also revealed a significant delay in the maturation of dorsal mLVs relative to the basal mLVs and cpLVs^15^, suggesting that the different networks of CNS lymphatics are phenotypically distinct from each other. Additionally, despite the capability of undergoing VEGFR3-dependent lymphangiogenesis after administration of VEGFC into the cisterna magna, dorsal mLVs do not undergo lymphangiogenesis during EAE^14,46^. One explanation is that cpLVs have greater access to pro-lymphangiogenic VEGFC molecules within the CNS. Anatomically, the dorsal mLVs are separated from the CSF-filled subarachnoid space by the most significant margin, with basal mLVs residing in much closer proximity to the SAS with seemingly greater access to CSF^16^. Here, we show cpLVs may have direct access to the SAS with gaps in E-Cadherin expression where cpLVs are present. Additionally, cpLVs have access to a CSF reservoir, express the water pore AQP-1, and dye infused into the CSF can be found within cpLVs. Heterogeneity in CSF access may reflect different levels of sampling from the CNS and therefore variation in their respected effects on disease states.

Another potential difference between the dorsal mLVs and cpLVs can be seen with inhibition studies. Photo-inhibition of specifically the dorsal mLVs using Visudyne during EAE results in decreased EAE severity^46^, suggesting a contribution of the dorsal mLVs to EAE pathology. In contrast, one of the phenotypic changes by cpLVs during EAE is their ability to bind and interact with dendritic cells to potentially engage in antigen transfer and tolerance through PDL-1. These findings suggest a role in antigen processing, antigen presentation, and immunoregulation which we plan to study in more detail in the future. Interestingly, administration of the VEGFR3 tyrosine kinase inhibitor MAZ51 once per day before the onset of clinical symptoms reduces EAE severity by causing both meningeal lymphatic regression and inhibition of lymphangiogenesis; however, MAZ51 administration after the onset of EAE clinical symptoms results in no change in EAE severity^9^. These data suggest that at later time points in the more chronic phase of EAE, the dorsal mLVs and lymphangiogenic cpLVs may have opposing roles in promoting autoimmunity or tolerance, respectively. This is also consistent with the hypothesized dual, stage-specific role of IFN-γ during EAE^38^, which also seems to regulate some aspects of the cpLEC phenotype. In addition to heterogeneity between lymphatic systems within EAE, there is also heterogeneity between diseases. For example, in the case of brain tumors or aging, administering VEGFC to promote the dorsal and cpLV function increases anti-cancer immunity^47,48^ and cognitive function^49^. Thus, there seems to be heterogeneity between the dorsal mLVs, basal mLVs, and cpLVs in terms of their phenotype, functionality, response to different diseases, and development, which should be considered in future studies.

Antigen processing, presentation, and archiving have been shown to occur in lymphatic endothelial cells in the lymph nodes, highlighting a relatively novel role of lymphatic endothelial cells as active players in regulating immunity through interactions with dendritic cells and T cells^24,25,26,27,28^. In this study, we highlight the potential ability of cpLECs upstream of the lymph nodes to uptake CNS-derived antigens and express MHC II, which correlates with their binding to inflammatory dendritic cells and a lesser extent CD4 T cells. The uptake of antigens and the binding of different subsets of T cells likely depends on the model of neuroinflammation. Here, we used an EAE model in C57BL/6 mice in which antigen processing and presentation are mediated by dendritic cells^50^, followed by predominantly a CD4 T cell response^51^ against a myelin-specific self-antigen, in this case, MOG_35-55_. This is consistent with the cpLEC phenotype observed in this study in which there is an uptake of a CNS-derived oligodendrocyte-specific antigen along with MHC II expression, dendritic cell binding, and CD4 T cell binding. While CD4 T cells have been shown to migrate to lymphatics in a CCR7-dependent manner^46^, here we show an approximately 10-fold increase in dendritic cell to cpLECs binding relative to CD4 T cells and hypothesize that dendritic cells may remain the predominant cell type to bind to cpLECs during neuroinflammation due to their roles as professional antigen-presenting cells and ability to migrate to lymphatics in a CCL21-CCR7 dependent manner^9,31^. Future studies are needed to assess the precise timing of dendritic cell and CD4 T cell binding to cpLECs and the relative contributions to the different types of cellular crosstalk that are likely occurring in this region.

EAE induces a strong Th1/Th17 cell response, characterized by several cytokines, including IFN-γ. Although traditionally a pro-inflammatory cytokine, IFN-γ deficient mice have exacerbated EAE, suggesting a tolerogenic role of IFN-γ in regulating autoimmunity^40^. Our scRNAseq data highlighted a response to IFN-γ signaling within cluster 2, which consisted primarily of EAE cpLECs that were phenotypically unique compared to steady-state cells from cluster 1. Here, we demonstrate that IFN-γ is required for the upregulation of PDL-1 by cpLECs during EAE to potentially engage in tolerogenic crosstalk with CD4 T cells, which has only been shown to occur by lymph node LECs^31^. The mechanism by which cribriform plate LECs capture CNS antigens and interact with leukocytes and the long-term impact of LEC-DC crosstalk is the focus of future studies. Interestingly, visualizing cell trajectories also revealed that genes involved in the regulation of leukocyte activation such as PDL-1 seem to be enriched at a later point in pseudotime relative to genes involved in response to IFN-γ **(Supplementary Figure 3)**, which is in line with the observation that PDL-1 expression by both dendritic cells and cpLECs is regulated by IFN-γ **(Figure 4)**. In addition to PDL-1, IFN-γ also regulates Podoplanin expression, which has been shown to be involved in LEC development and the recruitment of dendritic cells^34,38^, which would coincide with our data showing a significant increase in the number of dendritic cells binding to cpLECs. Interestingly, lymphangiogenesis is not dependent upon IFN-γ signaling, as IFN-γ KO mice still undergo lymphangiogenesis **(Supplementary Figure 7)**. Although we can rule IFN-γ out as a driver of lymphangiogenesis, its role in the regulation of Podoplanin and PDL-1 expression highlight its importance for regulating other potential functions of lymphangiogenic cpLVs, and future studies are needed to elucidate the roles of other factors such as Type I interferons in regulating cpLEC phenotype^25^.

In conclusion, we have, for the first time, characterized cpLECs in-depth with scRNAseq in both steady-state and EAE to identify novel functions of cpLECs in the regulation of neuroinflammation. These data not only validate our initial findings of lymphangiogenesis by the proliferation of pre-existing LECs but show lymphangiogenic cpLECs undergo other functional changes, including leukocyte crosstalk and increased CSF sampling, some of which as a response to the IFN-γ-heavy microenvironment. Many of these functions, including antigen capture, antigen archiving, and PDL-1-PD-1 dependent T cell tolerance, have only recently been attributed to LECs in the lymph nodes, suggesting cpLECs upstream of the lymph nodes act as key players in regulating adaptive immunity. In this study, we provide new information regarding the complexity of CNS lymphatics and propose a balance of driving immunosurveillance through CSF sampling and antigen drainage with tolerance through ligands such as PDL-1 by cpLECs during neuroinflammation. This likely occurs alongside other CNS lymphatics such as the dorsal mLVs and basal mLVs, which seem to be heterogeneous and distinct from each other in terms of their functional roles during neuroinflammation and response to different diseases. Neuroinflammation not only expands the draining cpLVs but also changes the function of cribriform plate lymphatics, which may be important in the pathogenesis of neuroinflammatory diseases.

## Acknowledgments

We thank Khen Macvilay for his expertise in flow cytometry, Laura Schmitt-Brunold for her expertise in molecular biology, and all our laboratory members for insightful discussions and constructive criticisms of this work. We would like to thank Lauren Nettenstrom and Abigale Bleil at the UW Flow Core Facility for their assistance and expertise in FACS sorting. Additionally, we would like to thank Tyler Duellman and Sandra Splinter BonDurant for their expertise and assistance in scRNAseq. We would also like to thank Beth Rauch and Elizabeth Meyerand at the UW Small Animal Imaging Facility (SAIF) supported by the UWCCC grant P30 CA014520 to use its facilities and services. This work was supported by the National Institutes of Health grants NS10847 and NS103506 awarded to ZF, HL128778 awarded to MS, the Neuroscience Training Program T32-GM007507 to MH and CL, and the 10x Genomics Pilot Award AAD7125 awarded to ZF. MS and ZF are co-senior authors.

## Contributions

M.H.^1^, M.S., and Z.F. conceptualized the experiments and reviewed, revised the manuscript. M.H.^1^ performed the experiments, generated the figures, analyzed the data, and wrote the manuscript. A.M. performed the analysis and assisted in generating figures for the scRNAseq data. Y.H.C. assisted with the MRI experiments. C.L. assisted with the FACS for scRNAseq. C.L. and M.H.^3^ performed the IV CD45 timing experiments. M.S. and Z.F. are co-senior authors. All authors edited the manuscript.

## Methods

### Animals

Female C57BL/6J wild-type (stock #: 000664) and IFN-γ^-/-^ mice (stock #: 002287) were purchased from Jackson Laboratories. CD11c-eYFP transgenic reporter mice were a generous gift from Dr. Michel C. Nussenzweig at Rockefeller University. CNP-Cre transgenic mice were a generous gift from Brian Popco at the University of Chicago. pZ/EG-OP OVA_257-264_-OVA_323-339_ (OVA ^fl/fl^ mice) were generated by our lab as previously described in the C57BL/6 background^7^. OVA^fl/fl^ mice were crossed with CNP-Cre mice to generate CNP-OVA transgenic mice that express OVA^257-264^ and OVA^323-339^ fused to GFP under the CNS oligodendrocyte-specific CNPase promoter^7,9^. Eight to twelve-week old female mice were used for all EAE experiments along with the appropriate age and sex matched controls. All experiments were conducted in accordance with guidelines from the National Institutes of Health and the University of Wisconsin – Madison Institutional Animal Care and Use Committee.

### EAE Induction

EAE was induced in 8 – 12-week-old female mice by subcutaneous immunization with 100 ug of MOG_35-55_ emulsified in Complete Freund’s Adjuvant (CFA) between the shoulder blades. 200 ng of Pertussis Toxin (PTX) was injected intraperitoneally at 0 d.p.i. and 2 d.p.i. The onset of clinical scores was observed between day 8 and 12 post-immunization, and were assessed daily as follows: 0, no clinical symptoms; 1, limp/flaccid tail; 2, partial hind limb paralysis; 3, complete hind limb paralysis; 4, quadriplegia; 5, moribund. Intermediate scores were also given for the appropriate symptoms.

### Magnetic Resonance Imaging

MRI experiments were done using a 4.7T small animal MRI (Agilent Technologies Inc., Santa Clara, CA, USA) and acquired using VnmrJ (Agilent Technologies). After scout scans, isotropic 3D T1-weighted scans were used to detect gadolinium using the following parameters: TR = 9.3 ms; TE = 4.7 ms; Flip Angle = 20 degrees; Field of View = 40×20×20 mm; Resolution = 256×128×128: Averages = 4; Voxel Size ≈ 156 μm^3^. These resulted in a time scan of approximately 11 minutes. Animals were anesthetized using isoflurane administered through a nose-cone, and 10 μl of gadolinium was injected into the cisterna magna at a rate of 2 μl/minute. Respiratory rates and body temperature were monitored to ensure normal physiology. A baseline scan was acquired prior to gadolinium injection, and image processing was done using FIJI software.

### Histology

Mice were terminally anesthetized with isoflurane and transcardially perfused with 0.1M PBS followed by perfusion with 4% PFA in 0.1M PBS. Mice were then decapitated and the skin was removed from the whole heads using forceps and scissors to separate the skin from the muscle and ear canal. The whole heads were fixed in 4% PFA in 0.1M PBS overnight. The whole heads were then decalcified in 14% EDTA in 0.1M PBS for seven days, followed by cryoprotection in 30% sucrose in 0.1M PBS for three days. The EDTA was replaced with fresh 14% EDTA each day. The decalcified mouse heads were then embedded in Tissue-Tek OCT Compound, frozen on dry ice, then stored at −80°C. 60 μm thick frozen sections were obtained on a Leica CM1800 cryostat, mounted on Superfrost Plus microscope slides and stored at −80°C.

### Immunohistochemistry

Sections were thawed at room temperature for 10 minutes, washed with 0.1M PBS for 10 minutes, and unspecific binding was blocked with 10% BSA with 0.1% Triton-X in 0.1M PBS for 60 minutes. Sections were then incubated with the appropriate primary antibodies in 1% BSA and 0.1% Triton-X in 0.1M PBS at 4C overnight in a humidified chamber. The following antibodies were used for immunohistochemistry: Podoplanin PE (eBioscience, Catalog #: 12-5381-80), CD31 Alexa647 (BD Biosciences, Catalog #: 563608), Lyve-1 eFluor660 (Thermo Fisher Scientific, Catalog #: 50-0443-80), rabbit anti-GFP unconjugated (Novus Biologicals, Catalog #: NB100-1614-0.02ml), CNPase Alexa647 (Biolegend, Catalog #: 836407), MHC II eFluor450 (eBioscience, Catalog #: 48-5321-80), CD11c Alexa488 (Thermo Fisher Scientific, Catalog #: 53-0114-80), CD4 PE (BD Pharmingen, Catalog #: 553049), Goat anti-E-Cadherin unconjugated (R&D Systems, Catalog #: AF748), and rabbit anti-AQP-1 unconjugated (EMD Millipore, Catalog #: AB2219). Sections were then washed three times with 0.1M PBS for 10 minutes each, then incubated with the appropriate secondary antibodies in 1% BSA and 0.1% Triton-X in 0.1M PBS at room temperature for 120 minutes. The following secondary antibodies were used: Donkey anti-Chicken Alexa488 (Invitrogen, Catalog #: A11039), Donkey anti-Chicken Alexa647 (Invitrogen, Catalog #: A21449), Donkey anti-Goat Alexa568 (Invitrogen, Catalog #: A11057), and Donkey anti-rabbit Alexa568 (Invitrogen, Catalog #: A10042). Sections were then washed three times with 0.1M PBS for 10 minutes each, mounted with Prolong Gold mounting medium with DAPI, and images acquired using an Olympus Fluoview FV1200 confocal microscope. The brightness/contrast of each image was applied equally across the entire image and equally across all images and analyzed using either FIJI or IMARIS software.

### Single Cell Suspension

Mice were terminally anesthetized with isoflurane and transcardially perfused with PBS. Mice were then decapitated, and the skin was cut dorsal to the midline of the skullcap rostrally to expose the skullcap. The skullcap was then removed along with the brain after separation from the olfactory bulbs. The cribriform plate and its associated tissues, which included the olfactory bulbs, were dissected out and placed in a 70-micron strainer submerged in RPMI-1640 in a non-tissue culture treated petri dish. The tissues were then mechanically dissociated by pushing the tissue through the strainer using a syringe plunger. The mechanically dissociated cells were then spun down, washed, and resuspended in FACS buffer for staining.

### Flow Cytometry

After resuspension of mechanically dissociated cells in fluorescence-activated cell sorting (FACS) buffer (pH = 7.4, 0.1M PBS, 1 mM EDTA, 1% BSA), the cells underwent staining with conjugated antibodies/dyes. Conjugated antibodies were diluted 1:200 in FACS buffer, and the following antibodies and dyes were used: Ghost UV450 (Tonbo Biosciences, Catalog #: 13-0868-T500) to visualize live/dead cells; CD31 Alexa647 (BD Biosciences, Catalog #: 563608), Podoplanin PE (eBioscience, Catalog #: 12-5381-82), and Lyve-1 Alexa488 (eBioscience, Catalog #: 53-0443-82) to visualize lymphatic endothelial cells; CD45 APC-Cy7 (Biolegend, Catalog #: 103116) or CD45 APC eFluor780 (eBioscience, Catalog #: 47-0451-80) to visualize leukocytes; CD11b PerCP-Cy5.5 (Biolegend, Catalog #: 101227) or CD11b PE-Cy5 (Biolegend, Catalog #: 101210) to visualize myeloid cells; CD11c BV421 (Biolegend, Catalog #: 117329) or CD11c FITC (Biolegend, Catalog #: 117305) to visualize dendritic cells; CD8 PE-Cy7 to visualize cytotoxic T cells (Biolegend, Catalog #: 100721); CD4 BUV496 to visualize T helper cells (BD Bioscience, Catalog #: 564667); B220 BV510 to visualize B cells (Biolegend, Catalog #: 103247); and PDL-1 PE-Cy7 to visualize the tolerogenic ligand PDL-1 (Biolegend, Catalog #: 124313). After staining for 30 minutes at 4°C, cells were washed three times in FACS buffer and processed using a BD LSR II. For the CD45 IV timing experiments, cells were processed with Cytek’s 3-laser Northern Lights.

### FACS Sorting

After generating a single cell suspension and staining for cpLECs with Ghost UV450 (Tonbo Biosciences, Catalog #: 13-0868-T500), CD31 Alexa 647 (BD Biosciences, Catalog #: 565608), Podoplanin PE (eBioscience, Catalog #: 12-5381-82), and CD45 APC-Cy7 (Biolegend, Catalog #: 103116) as previously described, the cells were sorted with a FACS Aria III with a nozzle size of 130 um at the UW Flow Core satellite facility in the UW Biotechnology Center. Cells were sorted into FACS buffer (1% BSA in PBS). A total of 25,102 cells from the control group and 53,224 cells from the experimental group post-sort were provided to the Biotechnology Core for scRNAseq.

### In vivo CD45 IV labeling

EAE was induced in C57BL/6 wild-type mice, and at peak clinical scores, EAE mice were intravenously (IV) injected with CD45.2 antibody conjugated to BV711 (Biolegend, Catalog #: 109847) 3 minutes before harvest to control for *ex vivo* unspecific binding of blood-derived cells to LECs and to visualize direct *in vivo* access between blood vessels and lymphatic vessels. Unperfused mice were used as positive controls to visualize the blood-derived binding of leukocytes to cpLECs. Perfused mice were used for all experiments. cpLECs were then processed for flow as described previously, and processed using Cytek’s 3-laser Northern Lights.

### scRNA Sequencing

Five healthy mice were pooled for the control group, and 5 EAE mice were pooled for the experimental EAE group. A single-cell suspension of the cribriform plate and its associated tissues were generated as previously described, and cpLECs were FACS sorted for Ghost^negative^ live cells, singlets, CD45^low/negative^, Podoplanin^+^, and CD31^+^. Sorted cpLECs were then provided to the Biotechnology Core facility at the University of Wisconsin Madison for single-cell sequencing using the 10x Genomics Chromium Single Cell Gene Expression Assay. A total of 25,102 cells from the control group and 53,224 cells from the experimental group post-sort were provided to the Biotechnology Core for scRNAseq. The target recovery rate was 3,000 cells with a targeted read depth of 7,500. Cells were loaded onto a Chromium Controller to generate a single cell + barcoded gel-bead emulsion from which Illumina-compatible library preps were generated and sequenced on the NovaSeq in collaboration with the University of Wisconsin Biotechnology Center (UWBC) DNA Sequencing Facility. Data were then analyzed using a custom-developed single-cell data analysis pipeline generated by the UWBC Bioinformatics Resource Center. The resulting data were then analyzed and explored using the Loupe Cell Browser software. Significantly differentially expressed genes were defined using an adjusted P-value <0.1 using the Benjamini-Hochberg correction for multiple tests, excluding features that had an average occurrence of less than one count per cell.

### Gene ontologies

Genes identified to be differentially expressed from each cluster were separately assessed by gene ontological analyses using R package *clusterProfiler*^52^. The genes that passed filtering during alignment were used as the background set during over-representation analyses. Significant gene ontological terms were identified using an adjusted P-value < 0.05. Bar plots, gene-concept network plots, and enrichment maps were visualized and generated using *clusterProfiler*.

### Single-cell trajectory analysis

R package *monocle3*^21,22,23^ was employed for pseudotime analysis. Feature, gene annotation, and cell annotation matrices were regenerated for cells stemming from clusters of interest using the *cellranger reanalyze* function. A cell dataset was generated using these matrices and preprocessed using 50 as the maximum number of dimensions. Dimensions were reduced using UMAP methodology^53^. Cells were clustered, and trajectories were learned using default parameters by *monocle3*. Cells were ordered, and pseudotime was calculated in reference to a node in the largest cell cluster. Expression data was extracted for genes of interest and plotted against estimated pseudotime, using a minimum expression cutoff of 5 reads for visualization purposes.

## References

1. Tammela, T. & Alitalo, K. Lymphangiogenesis: molecular mechanisms and future promise. Cell 140, 460–476 (2010).

2. Alitalo, K. The lymphatic vasculature in disease. Nat. Med. 17, 1371–1380 (2011).

3. Engelhardt, B., Vajkoczy, P. & Weller, R. O. The movers and shapers in immune privilege of the CNS. Nat. Immunol. 18, 123–131 (2017).

4. Weller, R. O. et al. Pathophysiology of the lymphatic drainage of the central nervous system: Implications for pathogenesis and therapy of multiple sclerosis. Pathophysiology 17, 295–306 (2010).

5. Meyer, C., Martin-Blondel, G. & Liblau, R. S. Endothelial cells and lymphatics at the interface between the immune and central nervous systems: implications for multiple sclerosis. Curr. Opin. Neurol. 30, 222–230 (2017).

6. Johnston, M. et al. Evidence of connections between cerebrospinal fluid and nasal lymphatic vessels in humans, non-human primates and other mammalian species. Cereb. Fluid Res. 1, 2 (2004).

7. Harris, M. G. et al. Immune privilege of the CNS is not the consequence of limited antigen sampling. Sci. Rep. 4, 4422 (2014).

8. Laman, J. D. & Weller, R. O. Drainage of cells and soluble antigen from the CNS to regional lymph nodes. J. Neuroimmune. Pharmacol. 8, 840–856 (2013).

9. Hsu, M., Rayasam, A., Kijak, J.A. et al. Neuroinflammation-induced lymphangiogenesis near the cribriform plate contributes to drainage of CNS-derived antigens and immune cells. Nat Commun 10, 229 (2019).

10. Cserr, H. F. & Knopf, P. M. Cervical lymphatics, the blood-brain barrier and the immunoreactivity of the brain: a new view. Immunol. Today 13, 507–512 (1992).

11. Kida, S., Pantazis, A. & Weller, R. O. CSF drains directly from the subarachnoid space into nasal lymphatics in the rat. Anatomy, histology and immunological significance. Neuropathol. Appl. Neurobiol. 19, 480–488 (1993).

12. Koh, L., Zakharov, A. & Johnston, M. Integration of the subarachnoid space and lymphatics: is it time to embrace a new concept of cerebrospinal fluid absorption? Cereb. Fluid Res. 2, 6 (2005).

13. Norwood JN, et al. Anatomical basis and physiological role of cerebrospinal fluid transport through the murine cribriform plate. Elife. 8:e44278 (2019).

14. Louveau, A. et al. Structural and functional features of central nervous system lymphatic vessels. Nature 523, 337–341 (2015).

15. Aspelund, A. et al. A dural lymphatic vascular system that drains brain interstitial fluid and macromolecules. J. Exp. Med. 212, 991–999 (2015).

16. Ahn, J.H., Cho, H., Kim, J. et al. Meningeal lymphatic vessels at the skull base drain cerebrospinal fluid. Nature 572, 62–66 (2019).

17. Weller, R.O., Sharp, M.M., Christodoulides, M. et al. The meninges as barriers and facilitators for the movement of fluid, cells and pathogens related to the rodent and human CNS. Acta Neuropathol 135, 363–385 (2018).

18. Maruyama, K. et al. Inflammation-induced lymphangiogenesis in the cornea arises from CD11b-positive macrophages. J. Clin. Invest. 115, 2363–2372 (2005).

19. Zumsteg, A. et al. Myeloid cells contribute to tumor lymphangiogenesis. PLoS ONE 4, e7067 (2009).

20. Kerjaschki, D. The crucial role of macrophages in lymphangiogenesis. J. Clin. Invest. 115, 2316–2319 (2005).

21. Trapnell C, Cacchiarelli D, Grimsby J, et al. The dynamics and regulators of cell fate decisions are revealed by pseudotemporal ordering of single cells. Nat Biotechnol. 2014;32(4):381–386.

22. Qiu, X., Hill, A., Packer, J. et al. Single-cell mRNA quantification and differential analysis with Census. Nat Methods 14, 309–315 (2017).

23. Qiu, X., Mao, Q., Tang, Y. et al. Reversed graph embedding resolves complex single-cell trajectories. Nat Methods 14, 979–982 (2017).

24. Tewalt EF, Cohen JN, Rouhani SJ, Guidi CJ, Qiao H, Fahl SP, et al. Lymphatic endothelial cells induce tolerance via PD-L1 and lack of costimulation leading to high-level PD-1 expression on CD8 T cells. Blood. (2012) 120:4772–82.

25. Lucas ED, Finlon JM, Burchill MA, McCarthy MK, Morrison TE, Colpitts T. MB, et al. Type 1 IFN and PD-L1 coordinate lymphatic endothelial cell expansion and contraction during an inflammatory immune response. J Immunol. (2018) 201:1735–47.

26. Rouhani, S., Eccles, J., Riccardi, P. et al. Roles of lymphatic endothelial cells expressing peripheral tissue antigens in CD4 T-cell tolerance induction. Nat Commun 6, 6771 (2015).

27. Lucas E.D., Tamburini B.A.J. Lymph node lymphatic endothelial cell expansion and contraction and the programming of the immune response. Front. Immunol. 2019;10:36.

28. Santambrogio L, Berendam SJ, Engelhard VH. The antigen processing and presentation machinery in lymphatic endothelial cells. Front Immunol. (2019) 10:1033

29. Bendall, S.C. Diamonds in the doublets. Nat Biotechnol 38, 559–561 (2020).

30. Giladi, A., Cohen, M., Medaglia, C. et al. Dissecting cellular crosstalk by sequencing physically interacting cells. Nat Biotechnol 38, 629–637 (2020).

31. Clarkson B.D., Héninger E., Harris M.G., Lee J., Sandor M., Fabry Z. (2012) Innate-Adaptive Crosstalk: How Dendritic Cells Shape Immune Responses in the CNS. In: Lambris J., Hajishengallis G. (eds) Current Topics in Innate Immunity II. Advances in Experimental Medicine and Biology, vol 946. Springer, New York, NY.

32. Tamburini, B., Burchill, M. & Kedl, R. Antigen capture and archiving by lymphatic endothelial cells following vaccination or viral infection. Nat Commun 5, 3989 (2014).

33. Marelli-Berg FM, Clement M, Mauro C, Caligiuri G. An immunologist’s guide to CD31 function in T-cells. J. Cell Sci. 2013;126:2343–2352.

34. Acton SE, Astarita J, Malhotra D, et al. Podoplanin-rich stromal networks induce dendritic cell motility via activation of C-type lectin receptor CLEC-2. Immunity 2012;37:276–89.

35. Johnson, L., Banerji, S., Lawrance, W. et al. Dendritic cells enter lymph vessels by hyaluronan-mediated docking to the endothelial receptor LYVE-1. Nat Immunol 18, 762–770 (2017)

36. Torzicky M. Viznerova P. Richter S. Strobl H. Scheinecker C. Foedinger D. Riedl E. Platelet endothelial cell adhesion molecule-1 (PECAM-1/CD31), CD99 are critical in lymphatic transmigration of human dendritic cells. J Invest Dermatol. 2012;132:1149–1157

37. Fletcher JM, Lalor SJ, Sweeney CM, Tubridy N, Mills KH. T cells in multiple sclerosis and experimental autoimmune encephalomyelitis. Clin Exp Immunol. (2010) 162:1–11.

38. Astarita JL, Acton SE, Turley SJ. Podoplanin: emerging functions in development, the immune system, and cancer. Front Immunol. 2012;3:283.

39. Lane R. S., Femel J., Breazeale A. P., Loo C. P., Thibault G., Kaempf A., et al. (2018). IFNgamma-activated dermal lymphatic vessels inhibit cytotoxic T cells in melanoma and inflamed skin. J. Exp. Med. 215, 3057–3074.

40. Arellano G, Ottum PA, Reyes LI, Burgos PI, Naves R. Stage-specific role of interferon-gamma in experimental autoimmune encephalomyelitis and multiple sclerosis. Front. Immunol. 2015;6:492.

41. Papadopoulos M. C., Verkman A. S. (2013). Aquaporin water channels in the nervous system. Nat. Rev. Neurosci. 14, 265–277.

42. Trillo-Contreras, J. L., Toledo-Aral, J. J., Echevarria, M. & Villadiego, J. AQP1 and AQP4 contribution to cerebrospinal fluid homeostasis. Cells 8. 10.3390/cells8020197 (2019)

43. Mestre H., Hablitz L.M., Xavier A.L., Feng W., Zou W., Pu T., Monai H., Murlidharan G., Castellanos Rivera R.M., Simon M.J., et al. 2018. Aquaporin-4-dependent glymphatic solute transport in the rodent brain. eLife. 7:e40070

44. Abbott NJ, Pizzo ME, Preston JE, Janigro D, Thorne RG. The role of brain barriers in fluid movement in the CNS: is there a ‘glymphatic’ system? Acta Neuropathol. 2018;135:387–407. doi: 10.1007/s00401-018-1812-4

45. Trillo-Contreras, J. L., Toledo-Aral, J. J., Echevarria, M. & Villadiego, J. AQP1 and AQP4 contribution to cerebrospinal fluid homeostasis. Cells 8. 10.3390/cells8020197 (2019)

46. Louveau A, et al. CNS lymphatic drainage and neuroinflammation are regulated by meningeal lymphatic vasculature. Nat. Neurosci. 2018;21:1380–1391.

47. Hu X, Deng Q, Ma L, et al. Meningeal lymphatic vessels regulate brain tumor drainage and immunity. Cell Res. 2020.

48. Song E., Mao T., Dong H., Boisserand L.S.B., Antila S., Bosenberg M., Alitalo K., Thomas J.L., Iwasaki A. VEGF-C-driven lymphatic drainage enables immunosurveillance of brain tumours. Nature. 2020;577:689–694.

49. Da Mesquita S, et al. Functional aspects of meningeal lymphatics in ageing and Alzheimer’s disease. Nature. 2018;560:185–191.

50. Greter M, Heppner FL, Lemos MP, et al. Dendritic cells permit immune invasion of the CNS in an animal model of multiple sclerosis. Nat Med. 2005;11(3):328–334.

51. Kuchroo, V. K., A. C. Anderson, H. Waldner, M. Munder, E. Bettelli, L. B. Nicholson. 2002. T cell response in experimental autoimmune encephalomyelitis (EAE): role of self and cross-reactive antigens in shaping, tuning, and regulating the autopathogenic T cell repertoire. Annu. Rev. Immunol. 20:101.

52. Yu G, Wang L, Han Y, He Q (2012). “clusterProfiler: an R package for comparing biological themes among gene clusters.” OMICS: A Journal of Integrative Biology, 16(5), 284–287.

53. McInnes L, Healy J, & Melville J. UMAP: Uniform Manifold Approximation and Projection for Dimension Reduction. arXiv:1802.03426

