## Supplementary Information for "Neuroinflammation alters the phenotype of lymphangiogenic vessels near the cribriform plate"

**Supplementary Figure 1: cpLVs undergo lymphangiogenesis during EAE.**

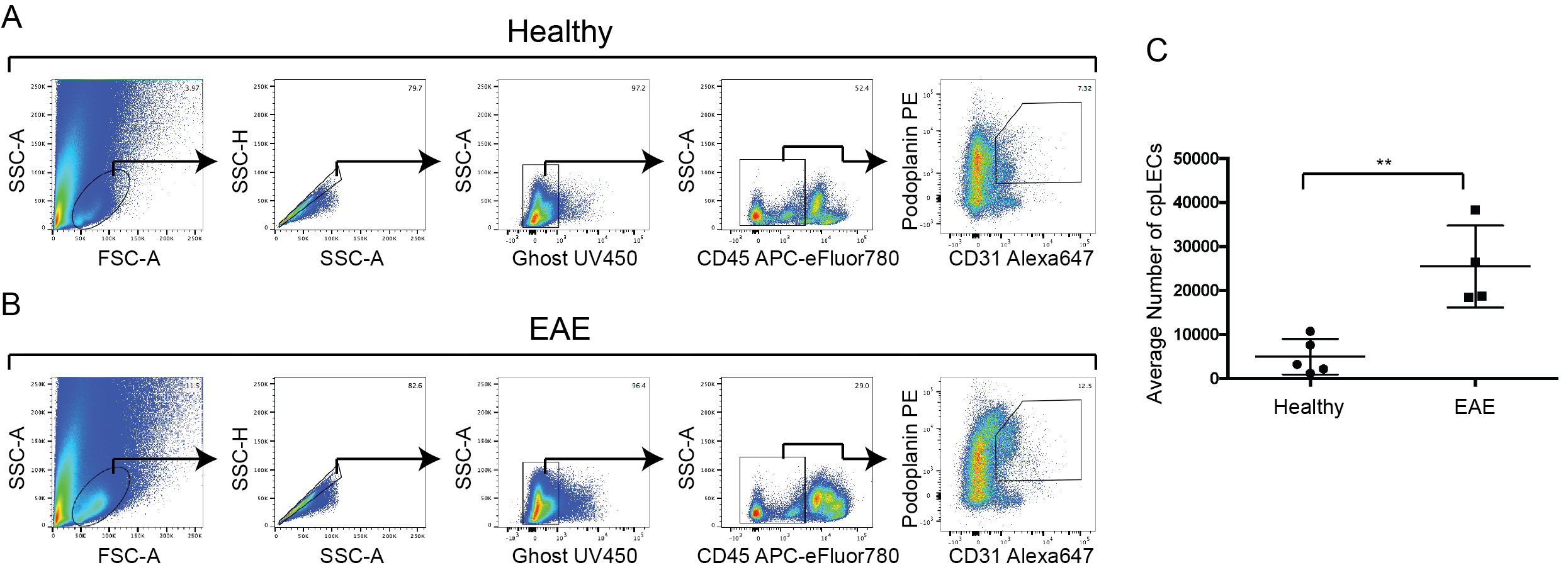

**Supplementary Figure 1: cpLVs undergo lymphangiogenesis during EAE.**

**(A):** Gating strategy from FACS sorting for scRNAseq of cpLECs.

**(B):** Gating strategy of the single cell suspension of cpLECs as shown in **(A)** between healthy and EAE mice. cpLECs were identified as Podoplanin^+^ CD31^+^ after excluding doublets, GhostUV450^+^ dead cells, and CD45^+^ leukocytes.

**(C):** Quantitation of the average cpLEC cell number by flow cytometry confirm lymphangiogenesis by cpLECs during EAE. *n* = 4 – 5 mice per group; data are represented as mean ± standard error of the mean, ***p* ≤ 0.01, unpaired Student’s t-test.

**Supplementary Figure 2: Volcano and CNET plots of scRNAseq data.**

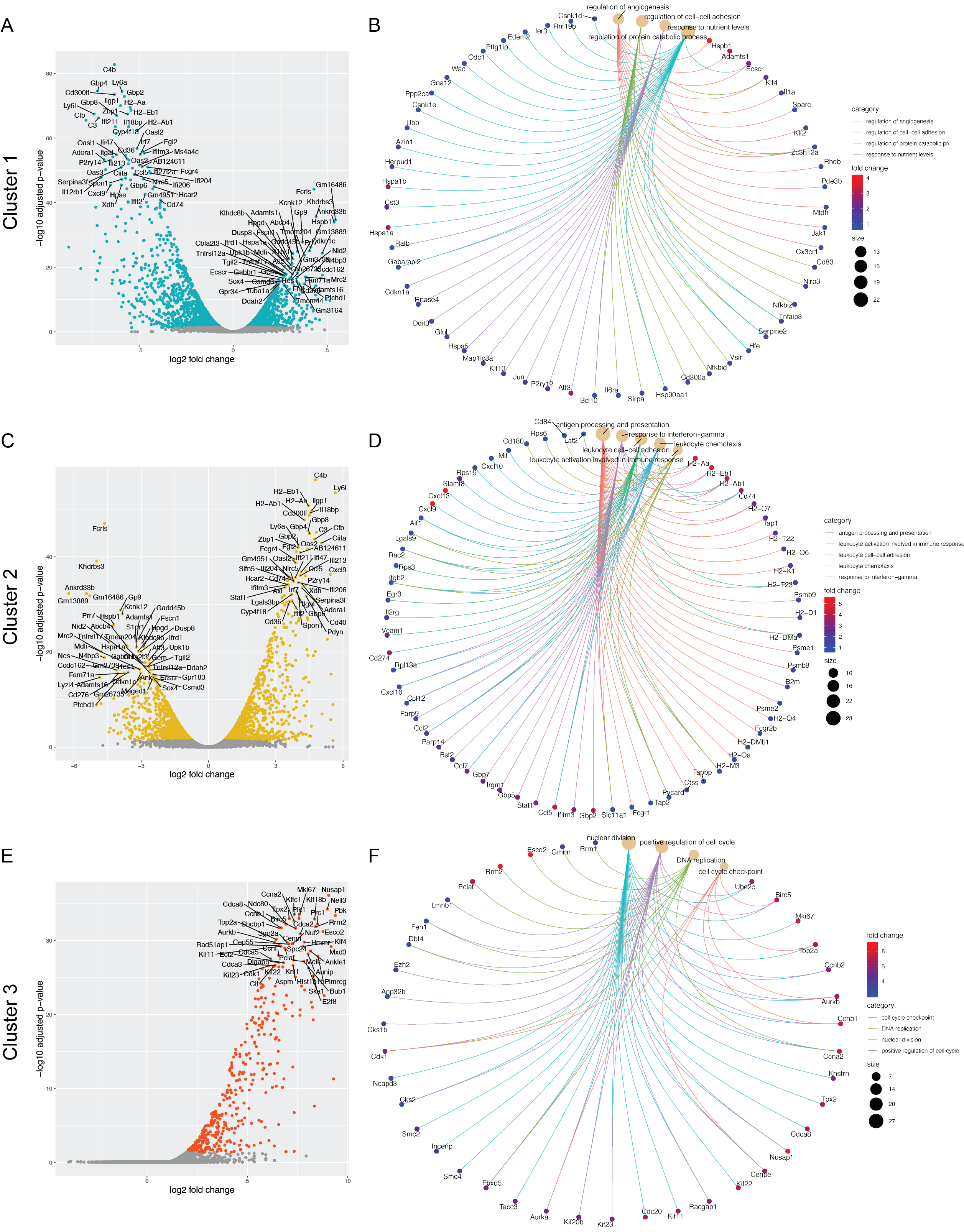

**Supplementary Figure 2: Volcano and Cnet plots of scRNAseq data.**

**(A):** Volcano plot showing the top 50 most up-regulated and down-regulated genes in Cluster 1.

**(B):** Cnet plot detailing the strength of association between representative GO enrichment terms for Cluster 1 for regulation of angiogenesis, regulation of protein catabolic process, regulation of cell-cell adhesion, and response to nutrient levels along with their associated genes.

**(C):** Volcano plot showing the top 50 most up-regulated and down-regulated genes in Cluster 2.

**(D):** Cnet plot detailing the strength of association between representative GO enrichment terms for Cluster 2 for antigen processing and presentation, response to interferon-gamma, leukocyte cell-cell adhesion, leukocyte chemotaxis, and leukocyte activation involved in immune response along with their associated genes.

**(E):** Volcano plot showing the top 50 most up-regulated genes in Cluster 3.

**(F):** Cnet plot detailing the strength of association between representative GO enrichment terms for Cluster 3 for nuclear division, positive regulation of cell cycle, DNA replication, and cell cycle checkpoint along with their associated genes.

**Supplementary Figure 3: Visualizing cpLEC trajectories.**

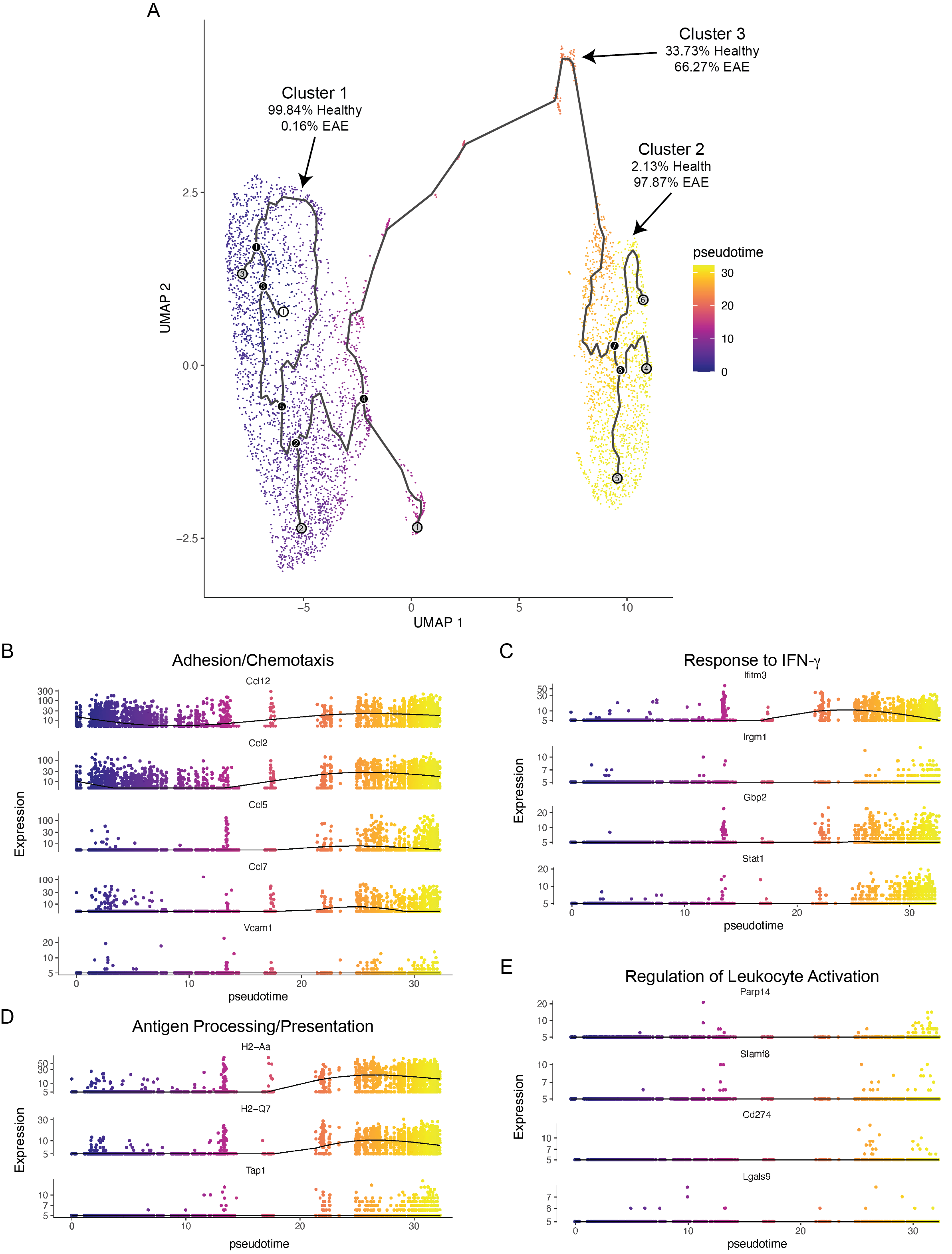

**Supplementary Figure 3: Visualizing cpLEC trajectories.**

**(A):** Single cell data was clustered using UMAP methodology and trajectories learned using default parameters by *monacle3*. Trajectories through pseudotime are shown for the three clusters; note that cluster 3 lies in between clusters 1 and 2 through pseudotime, and there are no direct connections between clusters 1 and 2.

**(B):** 5 representative genes associated with the GO enrichment term adhesion/chemotaxis that is enriched in cluster 2 are shown through pseudotime.

**(C):** 4 representative genes associated with the GO enrichment term response to IFN-γ that is enriched in cluster 2 are shown through pseudotime.

**(D):** 3 representative genes associated with the GO enrichment term antigen processing/presentation that is enriched in cluster 2 are shown through pseudotime.

**(E):** 4 representative genes associated with the GO enrichment term leukocyte activation that is enriched in cluster 2 are shown through pseudotime. Note that these genes tend to be up-regulated later through pseudotime relative to genes associated with the other 3 enrichment terms.

**Supplementary Figure 4: Leukocytes that bind to cpLECs do not come directly from blood.**

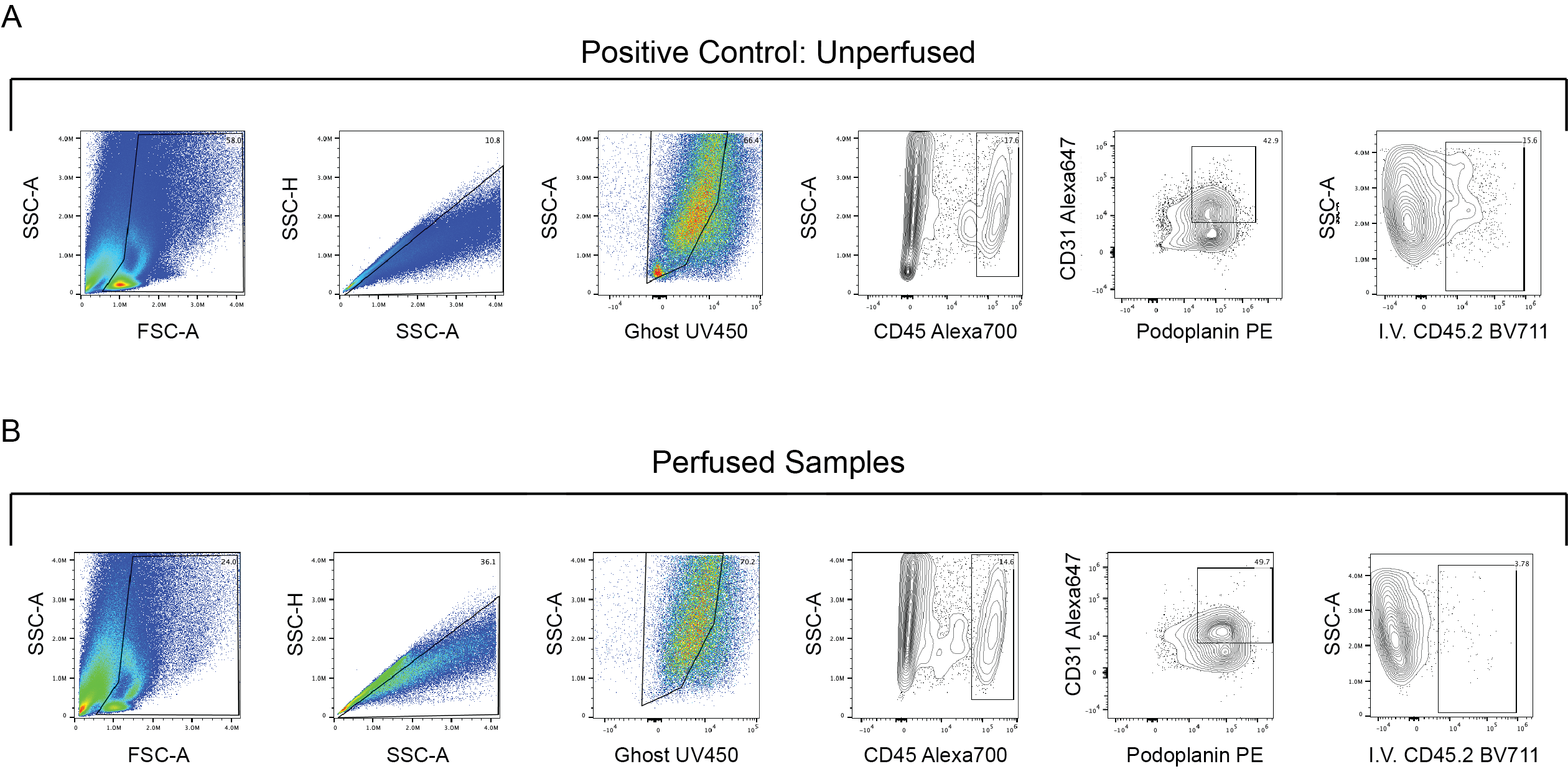

**Supplementary Figure 4: Leukocyte that bind to cpLECs do not come directly from blood.**

**(A – B):** Wild-type C57BL/6 mice were induced with EAE, and at peak EAE mice were intravenously (I.V.) injected with CD45.2 conjugated to BV711 antibody to label all blood-derived leukocytes. After allowing 3 minutes for the antibody to circulate and label blood-derived leukocytes, the mice were harvested for flow cytometry. To visualize blood-derived leukocyte that bind to cpLECs, unperfused **(A)** or perfused **(B)** mice were gated on live GhostUV450^-^ doublets, and leukocyte – cpLEC binding were visualized as CD45^+^ CD31^+^ Podoplanin^+^ **(A, B)**. Of the leukocytes bound to cpLECs in the unperfused control, approximately 16% were labeled with the 3-minute I.V. CD45.2 BV711 antibody **(A)**. Negligible amounts (< 4%) of the leukocytes bound to cpLECs in the perfused mice were labeled with the 3-minute I.V. CD45.2 BV711 antibody **(B)**.

**Supplementary Figure 5: EAE alters cpLV expression of CD31, PDPN, Lyve-1, and PDL-1.**

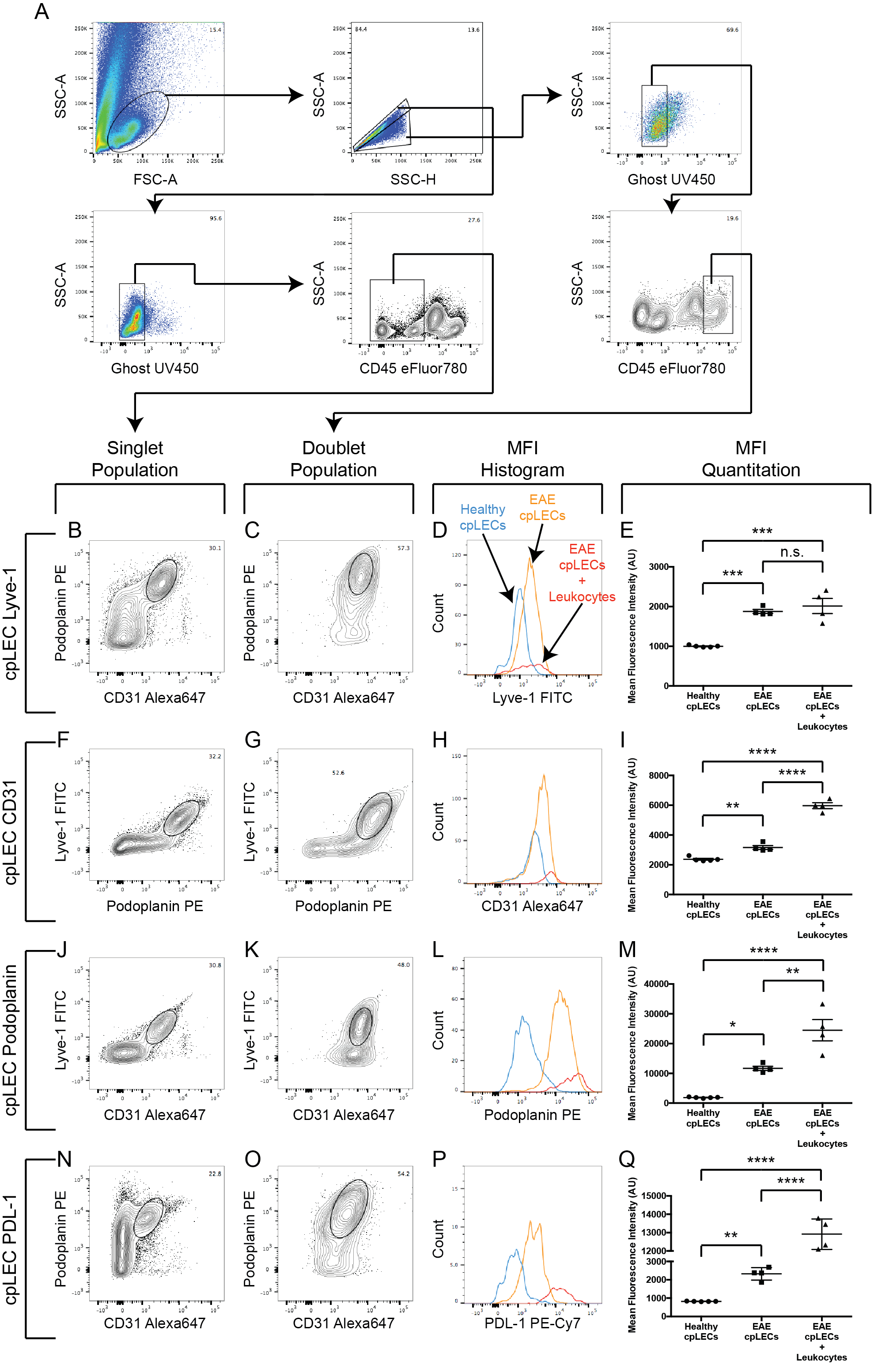

**Supplementary Figure 5: EAE alters cpLV expression of CD31, PDPN, Lyve-1, and PDL-1.**

**(A):** Gating strategy used to confirm the up-regulation of CD31, Podoplanin, Lyve-1, and PDL-1 at the protein level during EAE.

**(B – Q):** After gating for cpLECs as either Podoplanin^+^ CD31^+^ **(B – E, N – Q)**, Podoplanin^+^ Lyve-1^+^ **(F – I)**, or Lyve-1^+^ CD31^+^ **(J – M)**, the median fluorescence intensity (MFI) of Lyve-1 **(B – E)**, CD31 **(F – I)**, Podoplanin **(J – M)**, and PDL-1 **(N – Q)** by both singlet cpLECs and doublets in which a cpLEC is bound to a CD45^+^ leukocyte. *n* = 4 – 5 mice per group; data are represented as mean ± standard error of the mean, **p* ≤ 0.05, ***p* ≤ 0.01, ****p* ≤ 0.001, *****p* ≤ 0.001, one-way ANOVA with Tukeys post-hoc multiple comparisons test.

**Supplementary Figure 6: Increases in background MFI of doublets is negligible relative to protein expression.**

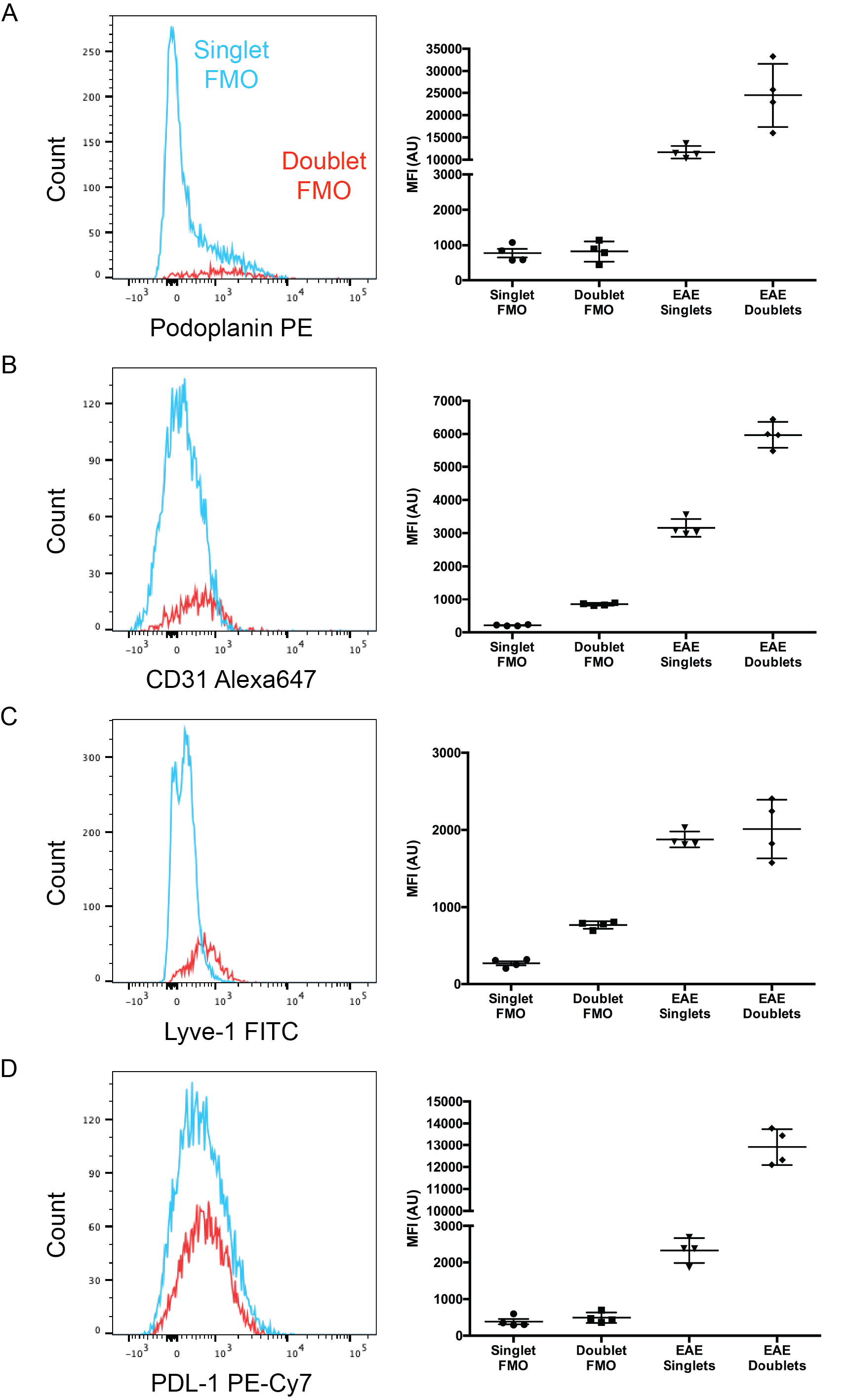

**Supplementary Figure 6: Increases in background MFI of doublets is negligible relative to protein expression.**

**(A – D):** FMO controls showing the increase in MFI of Podoplanin, CD31, Lyve-1, and PDL-1 of doublets relative to singlets due to background is negligible relative to actual protein expression.

**Supplementary Figure 7: cpLV lymphangiogenesis does not require IFN-γ signaling.**

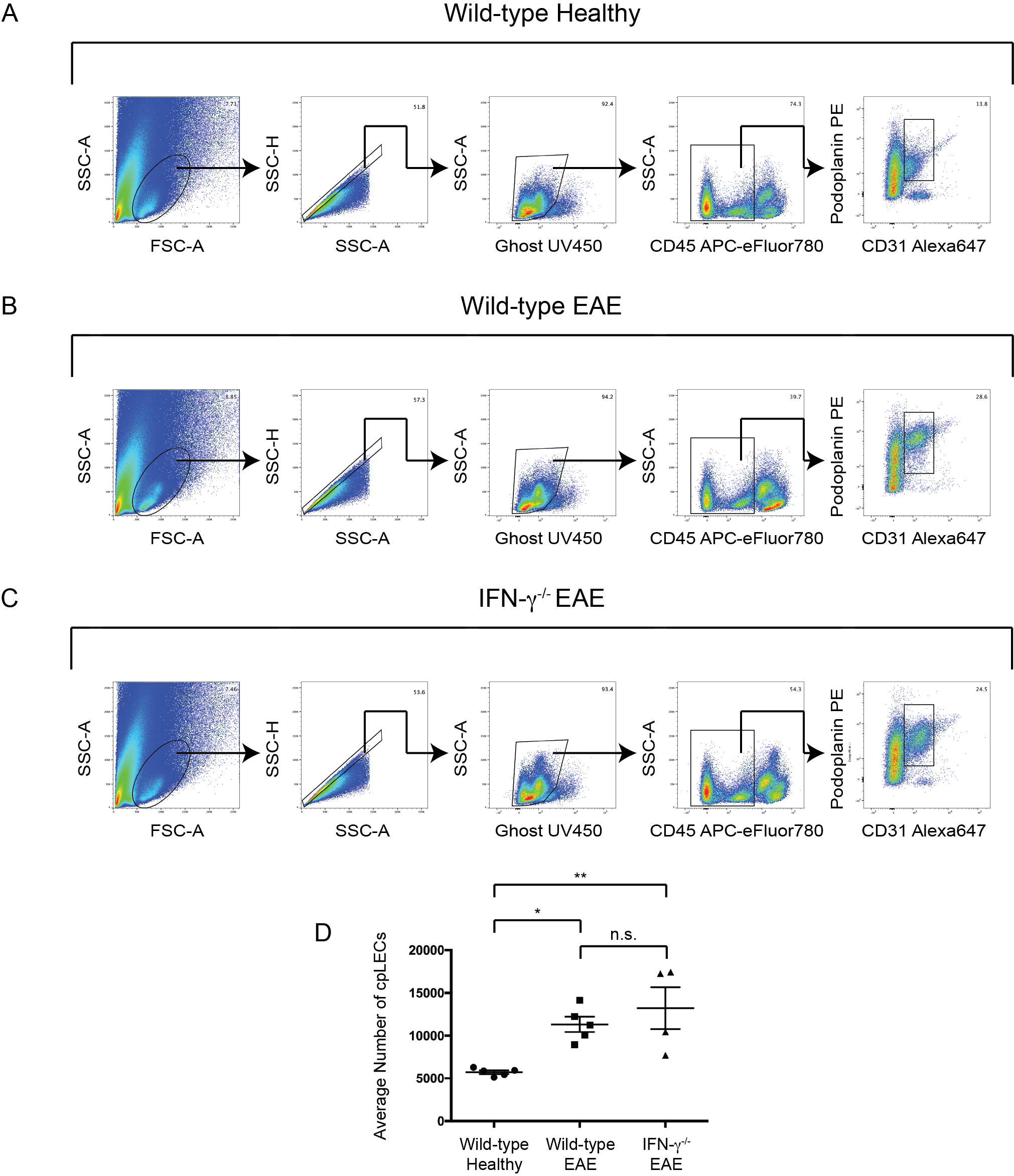

**Supplementary Figure 7: cpLV lymphangiogenesis does not require IFN-γ signaling.**

**(A – C):** Gating strategy for identifying cpLECs between wild-type healthy **(A)**, wild-type EAE **(B)**, and

IFN-γ^-/-^ EAE **(C)**.

**(D):** Quantitation of the average number of cpLECs between wild-type healthy **(A)**, wild-type EAE **(B)**, and

IFN-γ^-/-^ EAE **(C)** reveal no inhibition of lymphangiogenesis in IFN-γ deficient transgenic mice.  *n* = 4 – 5 mice per group; data are represented as mean ± standard error of the mean, **p* ≤ 0.05, ***p* ≤ 0.01, one-way ANOVA with Tukeys post-hoc multiple comparisons test.
